# Genome-wide modeling of DNA replication in space and time confirms the emergence of replication specific patterns in vivo in eukaryotes

**DOI:** 10.1101/2025.04.23.650322

**Authors:** Dario D’Asaro, Jean-Michel Arbona, Vinciane Piveteau, Aurèle Piazza, Cédric Vaillant, Daniel Jost

**Affiliations:** Laboratoire de Biologie et Modélisation de la Cellule École Normale Supérieure de Lyon, CNRS, UMR5239, Inserm U1293 Université Claude Bernard Lyon 1, 46 Allée d’Italie, 69007 Lyon, France; École Normale Supérieure de Lyon, CNRS, Laboratoire de Physique, 46 Allée d’Italie, 69007 Lyon, France

## Abstract

Although significant progress has been made on our understanding of DNA replication and spatial chromosome organization in eukaryotes, how they both interplay remains elusive. In particular, from the local structure of two diverging sister-forks to the higher-level organization of the replication machinery into nuclear domains, the mechanistic details of chromatin duplication in the 3D nuclear space remain debated. In this study, we use a computational model of the *Saccharomyces cerevisiae* genome to explore how replication influences chromatin folding. By integrating both a realistic description of the genome 3D architecture and 1D replication timing, simulations reveal that the colocalization of sister-forks produce a characteristic “fountain” pattern around early origins of replication. We confirm the presence of similar features *in vivo* in early S-phase with new Hi-C data in various conditions, showing that it is replication-dependent and cohesin-independent. At a larger scale, we show that the 3D genome leads to forks being highly enriched at one pole of the nucleus in early S-phase, before later redistributing more homogeneously, and may favor the higher-order clustering of forks into Replication Foci, as observed in earlier microscopy experiments. Additionally, replication causes temporary chromatin slowdown and reduced mobility due to fork passage and sister chromatid intertwining. Overall, our model offers new insights into the spatial and dynamic organization of chromatin during replication in eukaryotes.

## I. INTRODUCTION

Understanding how living cells duplicate, functionally reorganize and transmit their genomic information represents a fundamental challenge in molecular biology. In eukaryotes, such a process initiates at multiples genomic positions, the so-called origins of replication, and proceeds to the copy of the genome. Origin firing is hierarchical and finely regulated, creating distinctive and heterogeneous patterns of replication timing along the linear genome, known as the “replication timing program” [1, 2]. While traditionally investigated as a unidimensional process along the genome, advances in 3D genomics [3, 4, 5, 6] highlighted a strong correlation between the tridimensional genome organization and the unidimensional replication dynamics that occurs within the crowded environment of the nucleus [7, 8, 9]. However, establishing causal relationships between these phenomena remains challenging and our understanding of these crosstalks is limited. While, on the one hand, chromatin local architecture and accessibility likely regulate the Replication Timing Program [7, 8], on the other hand, the active progression of replication forks may also influence the spatial folding of chromosomes. For this reason, over the past decades, significant efforts have been devoted to describe the organization of replication forks in the 3D nuclear space [10, 11, 12, 13, 14, 15, 16].

During DNA replication, a first layer of organization lies in the 3D architecture of a single replication bubble (or replicon), a scale also relevant for prokaryotic genomes [17]. Currently, two models have been proposed to describe the relative 3D organization of the two diverging forks emanating from the same firing event, the so-called sister-forks or sister-replisomes [18]. The first model proposes that sister-forks remain in close spatial proximity as they progress in opposite directions along the genome, resulting in the effective extrusion of the two Sister Chromatids (SC) [19, 16, 13]. In the second “train-track” model [17, 20], sister-forks can instead diffuse independently from each other in the 3D space as replication proceeds. Despite various experimental evidence of the interdependent motion of sister-forks both in bacteria [21, 22] and eukaryotes [19, 16, 13], independently moving sister-forks have been also observed *in vivo* during bacterial replication [17, 23, 20] and *in vitro* in Xenopus egg extracts [20].

At a larger scale, early microscopy studies also reported the presence of large Replication Foci (RFi), each containing multiple forks, possibly aggregated through a phase separation mechanism [10, 12, 24, 11, 25]. While few observations made in budding yeast have supported this model [14], more recent super-resolution microscopy studies in mammals challenged it [26, 15], revealing that individual RFi can be further resolved into single replicons or forks. In this scenario, the previously observed, larger RFi may just arise from limited optical resolution combined with the simultaneous activation of multiple origins within the same 3D chromatin domain, without requiring specific, replication-dependent aggregating forces [26, 15].

Given such complex, multi-scale interplay between replication dynamics and 3D chromosome organization, quantitative approaches based on polymer simulations can be useful in testing different models and reconciling conflicting experimental observations. In the last couple of years, a few modeling studies have incorporated explicit replication into coarse-grained polymer frameworks to address these questions in bacteria [27, 28, 29, 30] and eukaryotes [19, 25, 31]. In bacteria, where replication starts from a single origin [27, 28, 29, 30], these studies suggest that the 3D genome architecture, driven by nucleoid confinement and SMC activity, coupled to replication, leads to the reliable segregation of SCs while promoting some degree of sister-fork colocalization. In eukaryotes, previous models [19, 25] explored idealized - toy - situations of eukaryotic-like replication with several replisomes working in parallel. In particular, our previous work highlighted how the local rearrangement of replication bubbles in S-phase may significantly impact the 3D structure and dynamics at different scales [19]. Forte *et al*. [25] characterize instead the energetic requirements to sustain sister-forks association via diffusing particles and their potential dynamics in the assembly of higher-order RFi-like structures. However, these approaches lack a realistic description of chromosome organization and origin firing that prevents quantitative analysis and comparison with experimental data.

To fill this gap, we develop a genome-wide 3D modeling of replicating chromosomes [19] of the budding yeast *Saccharomyces cerevisiae* for which the spatial architecture and replication are of minor complexity and have been extensively characterized separately. At the nuclear level, the *Saccharomyces cerevisiae* genome exhibits a polymer brush-like Rabl organization, where all 16 centromeres are attached to the Spindle Pole Body (SPB) via the mitotic spindle [32] while telomeres are anchored to the Nuclear Envelope (NE) [33, 34]. Opposite to the SPB lies the nucleolus, a defined subnuclear structure that isolates the ribosomal DNA repeats of chromosome 12 from the rest of the genome [35, 32]. Entering S-phase, yeast genome is duplicated starting from specific consensus sequences called Autonomous Replicating Sequences (ARS)[36, 37] and whose efficiencies are highly heterogeneous, leading to complex replication patterns [38, 39, 40, 41, 1, 42]. While G1 chromosomes lack of structural features such as TADs or loops [3], cohesin is gradually loaded on chromatin during S phase to mediate sister chromatids cohesion [43, 44, 45, 46, 47, 48, 49] and mitotic condensation via loop extrusion [50, 51, 52, 53, 54, 55]. To realistically simulate such a system, we couple a precise description of the hierarchical firing of yeast origins [56, 57] with a coarse-grained polymer model explicitly accounting for chain duplication [19] and for the budding yeast Rabl-organization [58, 59, 60, 61, 62]. This integrated approach enables us to examine the 3D features of *Saccharomyces cerevisiae* during S-phase across multiple scales. Taking advantage of the realistic 1D replication dynamics and genome-wide statistics, we first explore how the two proposed models of sister-fork association impact 3D chromosome structure at the replicon scale. Additionally, we compare our predictions with new *in vivo* Hi-C data from S-phase, which reveal distinct fountain-like patterns around early origins. These patterns are consistent with at least a fraction of sister-forks extruding the newly synthesized SCs during their progression.

Beyond the replicon scale, our framework allows to explore how the physical constraints imposed by the Rabl architecture shape the 3D distribution of forks across the nucleus, potentially affecting the detection of larger RFi containing multiple pairs of sister-forks.

Finally, we explore the dynamics of replicating chromosomes, providing quantitative insights on the potential slowdown in the diffusion of chromatin in the presence of active forks and intertwined structures.

## II. RESULTS

### A. Modeling replication in space and time

#### 1. A minimal polymer model recapitulates the yeast nuclear architecture in G1

In order to address the role of replication in driving the 3D Genome in S-phase, we first implement a minimal, quantitative genome-wide polymer model of the nuclear architecture of haploid yeast in G1. It aims at capturing the most fundamental structural properties of chromosome organization to serve as a “null” model.

Briefly (see Materials and Methods for details), similarly to previous studies [58, 59, 60, 61, 62], we model the full *Saccharomyces cerevisae* haploid genome by confining the 16 chromosomes into a spherical volume to mimic the

Nuclear Envelope (NE) (Fig. 1A, Fig. S1B,C). Each chromosome is modeled as a semi-flexible, self-avoiding chain, each monomer encompassing 1 kbp and being of size 20 nm. Specific features of the yeast typical Rabl-organization [35, 32] like the clustering of centromeres at the spindle pole body (SPB) and the tethering of telomeres at NE are recovered by adding physical constraints on the corresponding specific regions of each chromosome (Fig. 1E, Fig. S1B). Finally, we include a minimal description of the nucleolus, that encompasses the Mbp-long rDNA region present on chromosome 12, by including a barrier (“nucleolus wall”) within the nuclear sphere to mimic the 3D space inaccessible to the rest of DNA and by splitting chromosome 12 into two separate chains, each including a rDNA boundary which is tethered to the nucleolus wall (Fig. 1E, Fig. S1C).

**FIG. 1:**
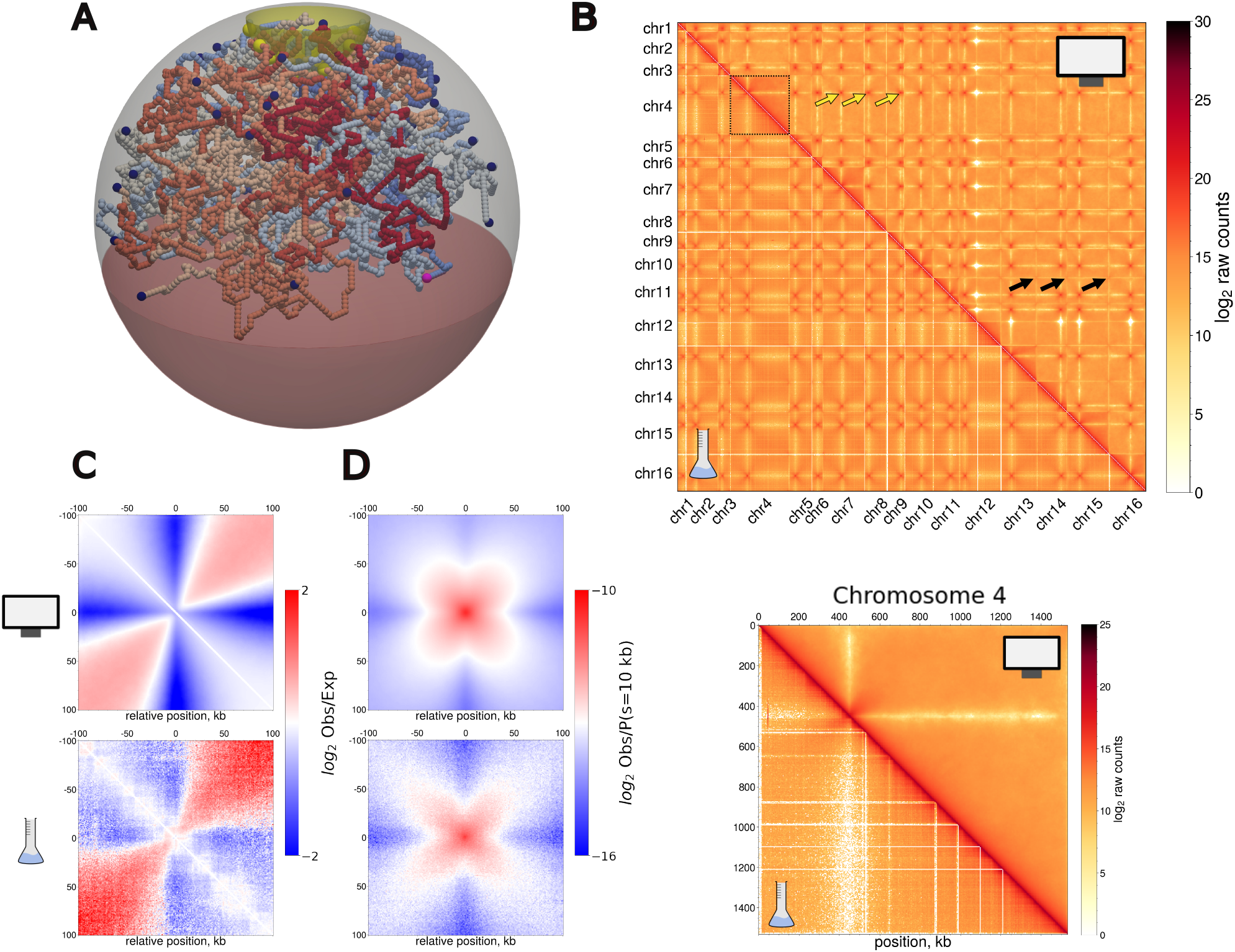
Genome-wide modeling of *Saccharomyces cerevisae* haploid genome in G1. A) Snapshot of G1-like simulations. Each color indicates a different polymer chain. Centromeres, telomeres and rDNA boundaries are highlighted with larger beads in yellow, dark blue and pink respectively. The red region within the sphere indicates the nucleolus and is inaccessible to other monomers. (B) Top: simulated (upper triangle) and *in vivo* (lower triangle) Hi-C maps (raw contacts) of the whole genome at a resolution of 16 kb for *r*_*c*_ = 80 nm. Yellow and black arrows indicate inter-centromeres and inter-telomeres contacts respectively. The dashed square on the diagonal highlights chromosome 4, plotted in the bottom panel at 1 kb resolution. (C) Simulated (Top) and experimental (Bottom) on-diagonal aggregate plots (Observed/Expected) around yeast’s 16 centromeres. (D) Simulated (Top) and experimental (Bottom) off-diagonal inter-chromosomal aggregate plots between all centromeres pairs. The signal is normalized by the *P*(*s* = 10*kbp*).

As highlighted previously [63, 60, 61, 64], such a minimal model is able to capture quantitatively the typical polymer-brush-like architecture observed experimentally with an overall Pearson correlation of 0.86 between simulated and experimental Hi-C maps (Fig. 1). In particular, at the genome scale, we observe a high frequency of contacts between centromeres (Fig. 1B, yellow arrows). This predicted contatct enrichment arises from the imposed colocalization of centromeres around the SPB (Fig. 1C) and resembles the experimental observations. Similarly, the focal telomere-telomere contacts observed in Hi-C are captured (Fig. 1B, black arrows; Fig.1D), without imposing any attractive energy between telomeres, just by imposing their confinement at NE.

Besides the overall nuclear organization, our model also quantitatively reproduces the intra-chromosome architectures in G1 (Fig. 1B, S1D and S2) with an average Pearson correlation between experimental and simulated HiC maps of 0.94 at the single chromosome level. Thanks to the imposed brush architecture, the model predictions match the enrichment of contacts between the arms of the same chromosome observed experimentally around centromeres (Fig. 1C) and at long-ranger range (Fig. S2C).

In summary, our minimal “null” G1 model captures the main features of budding yeast 3D nuclear architecture, allowing us to investigate its interplay with explicit chromosome replication.

#### 2. Modeling replication dynamics and explicit chain duplication during S-phase

To model the yeast 3D genome during S-phase, we include explicit polymer duplication to simulate the progressive synthesis of two sister chromatids (SCs) from each “maternal” - G1 - chromosome (Fig.2A). This is achieved by employing a specialized class of polymers that self-replicate starting from predefined genomic positions, the origins of replication [19]. Locally, origin firing consists of adding a new monomer and connecting it to the maternal chain, leading to the formation of a replication bubble (Fig.2C,D, see Materials and Methods for details). The monomers connected at the extremities of this bubble are the replication forks (green monomers in Fig.2C). Further duplication of such triple connectivity accounts for fork progression along the genome which we assume to be stochastic and happening at constant average speed (*v* = 2.2 kb/min [38]). In such a formalism, multiple growing bubbles can be easily introduced and tracked, with converging bubbles being merged together into larger loops (Fig.2A).

In one simulated trajectory, the dynamics of chain duplication thus depends on the set of origins that will be fired and their timing of firing. To account quantitatively for the correct firing dynamics in yeast, we couple our 3D framework with state-of-the-art modeling of 1D replication dynamics along the genome. Briefly, we integrate the stochastic model of eukaryotic DNA replication by Arbona *et al*. [56, 57] (Fig. 2A) where firing is modeled as a bimolecular reaction between a set of putative origins and a limiting number of firing factors (see Materials and Methods for details). The positions of potential origins (p-ori) are randomly selected among all the monomers with a probability proportional to the so-called “Initiation Probability Landscape Signal” (*IPLS*) (Fig. 2A, left and B, top, Fig. S3C) which is inferred from replication data using neural network [57]. Peaks in *IPLS* correspond to origins of replication and mostly coincide to the yeast autonomously replicating sequences (ARS) (Fig. S3) from the Oridb database [36].

**FIG. 2:**
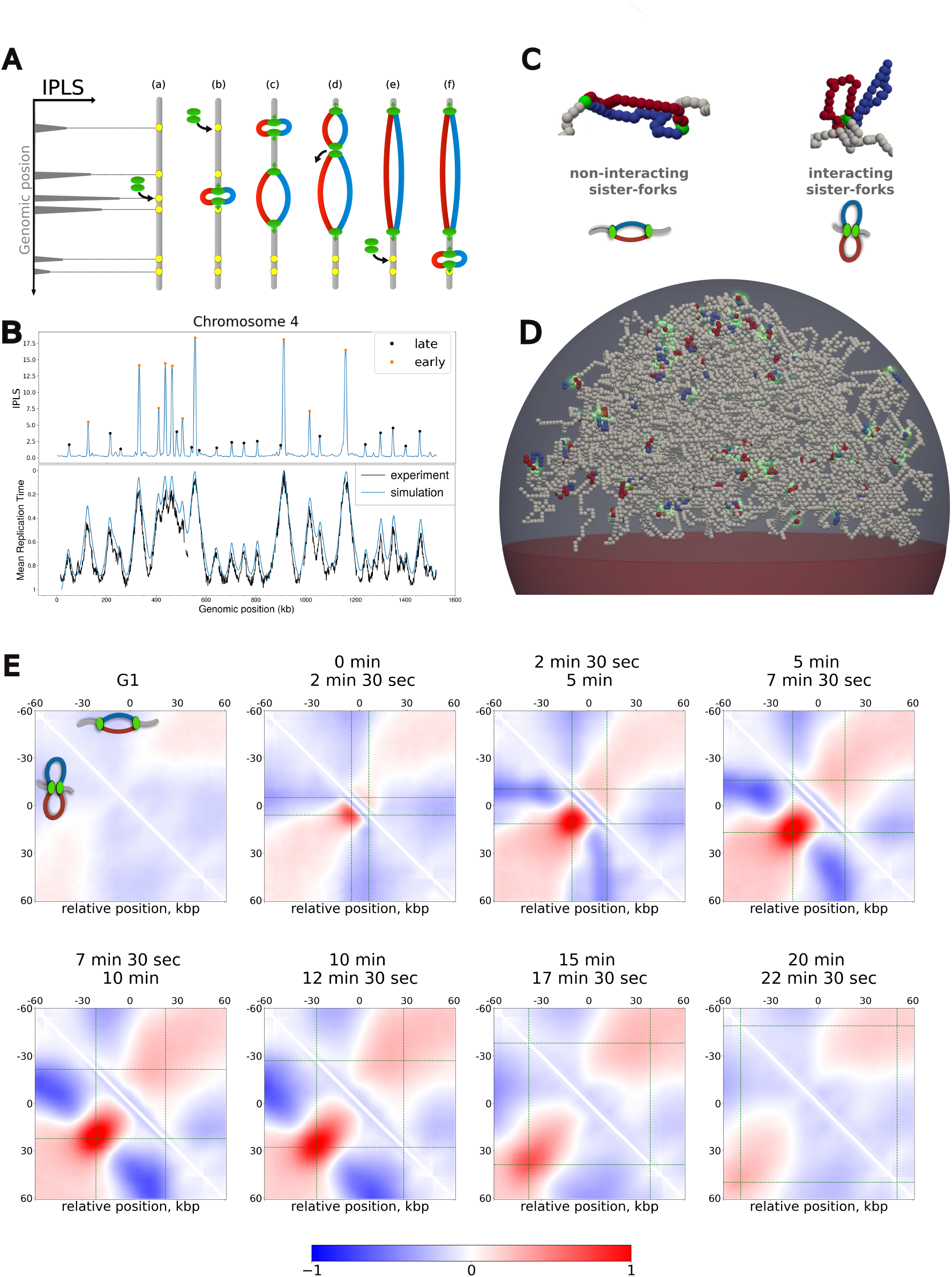
Spatio-temporal whole-genome model of yeast DNA replication. (A) Scheme of 1D replication model of Arbona *et al*. [56, 57]. Origins are drawn with replacement from the IPLS signal (left):(a,b) they “entrap” one firing factor leading to the opening of a replication bubble with two diverging forks; (c) Replication bubbles grow as forks progress, and new firing events might be hindered by the limited number of diffusing factors; (d) Converging bubbles (merging events) release a diffusing factor that can be employed to fire a new unreplicated origin(e,f). (B) IPLS of chromosome 4 (top) and corresponding predictions of the mean replication time (bottom, blue line) compared with the experimental MRT (from [65]) (bottom, black line). Orange and black circles over *IPLS* peaks indicate origin classified as early and late respectively (see Material and Methods). (C) The two investigated scenarios for the local sister-forks spatial organization with illustrative snapshots from simulations and corresponding scheme. Red and blue monomers correspond to the two replicated sister chromatids, light-gray to the unreplicated monomers. (D) Snapshot of a full genome simulation with chromosomes undergoing replication in the interacting sister-forks scenario. Green halos indicate replication forks location. (E) Average normalized (log_2_ Observed over Expected) contact maps around early replicating origins in the non-interacting (upper triangle) and interacting (lower triangle) scenarios at different times after the beginning of S-phase. Data are aggregated plots averaged over a time-window of 2.5 min. G1 corresponds to the unreplicated G1-like configurations. Green dashed lines indicate the expected average forks positioning at the corresponding time interval.

In particular, using *IPLS* (Fig. 2B top) and parameters inferred by Arbona *et al*. in [57], our model of replication dynamics well reproduces the MRT profiles observed in yeast all across the genome (Fig. 2B bottom, Fig. S4) and predicts a S-phase of duration ~ 30 minutes (Fig. S5) typical of yeast cell cycle [66]. Over the 726 inferred origins, we define sets of early and late origins (Fig.2B and S3, see Materials and Methods) depending on the peak height in *IPLS*, early origins (111 in total) having a very high probability to be fired at each S-phase, while late origins (615 in total) fired only occasionally.

Equipped with a modeling framework with realistic 1D replication dynamics and 3D chromatin folding within the nucleus, we investigate next how replication and chain duplication may impact genome organization during S-phase at different scales. In particular, we consider two scenarios (Fig. 2C): a first one where sister-forks evolve independently of each other (non-interacting scenario) and a second one where they are enforced to remain spatially colocalized (interacting scenario) until their replication bubble dissociates or merges with another bubble (see Materials and Methods). For each scenario, we simulate 300 S-phase trajectories, each initialized with a random G1-like configuration, and we track how the 3D chromosome organization evolves as a function of time after starting the replication process (Fig. 2D, Video S1).

### B. Replication impacts transiently the local chromatin organization around origins

We first focus on the local structural impacts of the formation of replication bubbles with or without interactions between sister-forks due to forks passage. From our simulations, we compute balanced Hi-C-like maps (see Materials and Methods) at different time points along the S-phase (Fig. S6).

#### 1. In silico prediction of chromatin fountains around early replicating origins

As early origins are fired frequently and more synchronously, we expect the surrounding genomic regions to be more exposed to any structural effects that fork passages may induce. In Fig. 2E, we plot the time evolution of the average contact enrichment around early origins in the interacting and non-interacting forks scenarios. We observe the emergence of typical, replication-dependent patterns that form very early in S-phase, evolve dynamically and finally progressively disappear as replication ends. Remarkably, both scenarios lead to qualitatively different features.

In the non-interacting case (upper triangles in Fig.2E), we detect a homogeneous enrichment of contacts in a band perpendicular to the main diagonal and centered around the origin whose width is related to the average replicon size at a given time around the origin (green dashed lines in Fig. 2E). This mild increase is related to the formation of replication bubbles around early origins [19]: the resulting ring-like topology of the polymer leads to an effective, entropically-driven compaction of the replicon compared to the rest of the unreplicated, linear polymer chain.

In the interacting scenario (lower triangles in Fig.2E), a stronger and more localized enrichment is clearly visible. Such an enrichment is maximal between the average positions of replication fork and progresses perpendicular to the main diagonal as replication proceeds. This typical signature of a fountain-like pattern, as also observed around cohesin loading sites [67, 68, 69, 70], results from the 3D colocalization of the two sister forks combined with their symmetrical progression along the genome around the origin with time. This leads to an effective loop extrusion mechanism [16, 19].

#### 2. Replication-dependent and cohesin-independent chromatin fountains are observed in vivo around early origins

We then investigate whether the predicted patterns can be detected in *in vivo* HiC data during S-phase. Although the presence of mild fountain-like patterns around origins in yeast was already reported in our previous study [19] using publicly available data in WT cells [55, 53, 46], it was not possible to directly establish a causal-effect relationship between these observations on HiC maps and the replication process as well as to rule out possible confounding effects such as the concomitant presence of extruding cohesins.

For this reason, we generate new HiC datasets during early S-phase (see Materials and Methods) in WT and mutant yeast strains to isolate the contribution of replication and decouple the emerging signal from concurrent biological processes: cells were arrested in G1 using alpha-factor and then released in S-phase for 20 minutes before fixation.

##### a. Enrichment around early origins in vivo is replication-dependent

In WT condition, analysis of the Hi-C data in early S-phase, shows the presence of a fountain-like pattern around early origins (Fig.3A Bottom left) with an enrichment around 10 - 30 kb. This structure appears specifically during S-phase as it is not visible in G1 (Fig.3A Top left) or G2/M-arrested (Fig.3A Top center) cells. Remarkably, the observed signal at this early-S timepoint appears significantly stronger than our previous report [19] using publicly-available Micro-C data [55].

To investigate the role of replication in the formation of the fountains, we perform similar experiments but in Cdc45-depleted cells using an auxin-degron. Cdc45 being a component of the replicative helicase CMG, it is essential for replication bubble opening at origins [71]. Cdc45-depleted cells thus progress towards G2/M but without performing DNA replication and in the absence of SC cohesion. In particular, it was observed that in this case chromosomes still engage in a mitotic-like compaction via cohesin-mediated loop extrusion activity[50, 52].

Measuring the Hi-C maps, as in the WT, 20 minutes after release from G1 and depletion in Cdc45 (Fig. S19C), we observe the complete loss of the fountain signal around early origins (Fig. 3A Top Right) after Cdc45 depletion. Overall, this result demonstrates that the fountain signal requires the presence of ongoing replication to be formed.

**FIG. 3:**
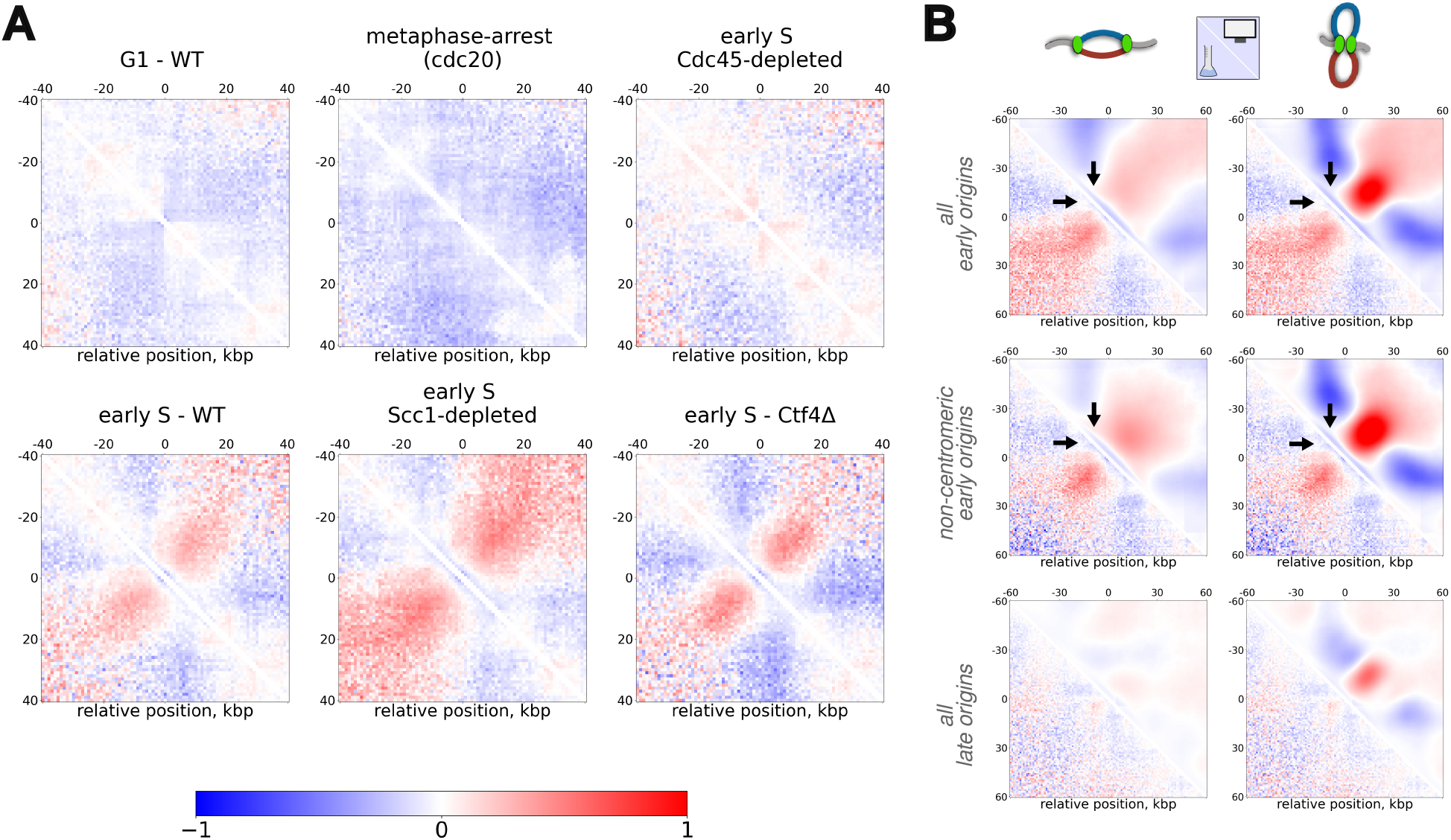
Comparison with *in vivo* data and forks spatial distribution. (A) Average normalized (log_2_ Observed over Expected) contact maps around early replicating origins for *in vivo* HiC data for different conditions. (B) Comparison between experimental aggregates of Scc1-depleted cells and simulation using the best matching time-interval from Fig. 2F (from *t* = 5 min to *t* = 7 min and 30, see Fig.S7A). Aggregates were computed around all early, only non-centromeric (centromeres at a distance *>* 50 kb) and all late origins for the two scenarios of sister-forks organization. Black arrows highlight the qualitative differences between the two simulated signals with a depletion and enrichment at the loop basis in the interacting and non-interacting scenario respectively.

##### b. Enrichment around early origins in vivo is cohesin-independent

Despite such findings, it is still unclear whether the emerging pattern is a direct consequence of the 3D organization of sister-forks, or driven by other concurring processes also happening during S-phase. In particular, in budding yeast, cohesin-mediated SCs cohesion and intra-chromatid loop-extrusion constitute two well-established processes that impact the 3D chromosome organization and that initiate during S-phase[43, 44, 45, 46, 47, 48, 49]. In particular, cohesins start extruding during early S leading to the formation of loops between specific genomic regions called Cohesin Associated Regions (CARs) [55] (Fig. S7B).

The early-S Cdc45-depleted and metaphase-arrested maps presented in Fig. 3A demonstrate that the fountains observed in WT early-S are not just a simple byproduct of some trivial correlations between CARs and early origins positioning along the genome, as a complete loss of fountains occurred in these cases while chromatin loops at CARS are still preserved (Fig. S7B). However, this doesn’t exclude that some cohesins might also be specifically recruited at replication forks and contribute to the signal.

To explore this hypothesis, we analyze early-S phase HiC map from cells partially depleted of Scc1 using an auxindegron (Fig. S19D), a cohesin subunit. We first verify that such a depletion reduce strongly the cohesin loop extruding activity and remove the loops between CARs in the Hi-C map, while their signature remains present in WT, Cdc45-depleted cells (Fig. S7B). Around early origin on the contrary, fountains are persistent to cohesin depletion (Fig. 3A Bottom Middle).

Overall, this suggests that the fountain signal does not depend on cohesin loading and loop extrusion activity.

##### c. Enrichment around early origins in vivo is Ctf4-independent

Given the experimental evidence of its ability to oligomerize *in vitro* [72], Ctf4, a component of the replisome, has been proposed as a possible mediator of sister-fork interactions. We thus perform Hi-C experiments as before but in Ctf4-KO cells that are still progressing through S-phase (Fig. S19E). In this case, we still observe loops around CARs (Fig. S7B) and more importantly, the characteristic enrichment around early origins is still present (Fig. 3A, Bottom Left), suggesting that Ctf4 might not be a driver of the fountain pattern.

#### 3. Comparisons with simulations suggest that sister-forks can interact in vivo

The Scc1-depleted maps discussed above offer the best comparison to our polymer model, which does not account for cohesin-mediated processes such as cohesion or loop-extrusion and only implements the formation and growth of replication bubbles.

To make the most meaningful comparison possible, we first have to infer the effective time in our simulations corresponding to the early-S stage in the experiments. To do so, we estimate and match the simulated and observed average numbers of Hi-C reads around early origins that reflects the local probability to have been replicated (Fig. S7A). We find a corresponding time interval in our simulations between 5 and 7.5 minutes (Fig. 2E, top right).

#### a. Around early replicating origins

We then compare the contact signals observed around early origins (Fig. 3B). Overall (Top), experimental data are qualitatively more similar to the fountains observed in the interacting fork scenario with common characteristic features: (1) a well-localized enrichment perpendicular to the main diagonal, corresponding to an average replicon size of 30 kbp, and (2) a depletion of the signal at the loop basis (black arrows in Fig.3B Top).

Since early origins are more densely located around centromeres, to avoid any possible structural artifacts related to the contact enrichment observed around centromeres (Fig.1C), we also compare specifically the signal for non-centromeric, early origins (Fig.3B Middle). They are characterized on average by greater inter-origin distances and thus are likely to have experienced less merging events between nearby replicons in early-S. *In vivo*, we still observed a fountain-like pattern which resembles also qualitatively more to the predictions of the interacting fork scenario.

Overall,the interacting fork predictions are thus more compatible with the fountain shape observed in the experimental data.

However, the strength of the enrichment signal is much stronger in the predictions than in the experiments (see also Fig. S7, S8). This may suggest an intermediate scenario where not all the sister-forks are paired. Indeed, our working hypothesis of strictly-bounded sister forks is likely to be a strong approximation of the real biological system as it has been experimentally observed that replication with independently moving forks is still viable [20].

We address this hypothesis by mixing contact maps of the interacting and non-interacting fork cases at different ratios (Fig. S7E). Interestingly, a small percentage of interacting forks signal (≥ 20%) is already sufficient to retain the localized enrichment perpendicular to the main diagonal (fountain pattern) while lowering the absolute value of the enrichment.

We also explore alternative confounding factors that may impact the predicted fountain strengths. While our model captures the experimental 1D dynamics and hierarchical firing of origins, all of the simulated trajectories are perfectly synchronized and enter replication at the same time, which is likely not the case in the experiments as the beginning of the S-phase in cycling cells or in cells released from G1 after synchronization is likely to be more heterogeneous [55, 51]. If we mix predictions for early-S with some for G1 (Fig. S7D), we observe also a lowering of the signal around early origins. Interestingly, the fountain pattern is very robust even at strong dilution suggesting that such replication-dependent signals may also be captured in a non-ideally synchronized system.

Finally, we investigate the impact of model parameters used in estimating the *in silico* contact maps: the radius of capture (Fig. S7G) which is the maximal distance between two monomers used to define a contact; and the length of the time intervals (Fig. S7F) used to compute maps. While these parameters have a quantitative impact on the enrichment, they both maintain the overall qualitative difference between the two fork scenarios.

Overall, experimental data around early origins are consistent with the presence of a significant populations of interacting sister-forks *in vivo*.

##### b. Around late replicating origins

We then investigate if our model also well-captures the signal around late replicating origins (Fig. 2B, Top). In the *in vivo* Scc1-depleted maps, we record only a faint signal around late origins, both in terms of size and strength (Fig. 2B Bottom, lower triangle). Predictions in both sister-fork scenarios show strong differences. In the interacting fork case, we continue to record a mild fountain-like pattern which is not present in the experiment (Fig. 2B Right). On the other hand, in the non-interacting case (Fig. 2B Left), only a faint signal is present, resembling more the experimental observation. Mixed scenarios with a majority of non-interacting cells are also consistent with the experiments (Fig. S9).

Overall, this suggests that interactions between sister-forks is less likely when fired from a late origin.

### C. The spatial organization of replication forks inside the nucleus is dynamic and heterogeneous

Thanks to our genome-wide modeling of the Rabl-like yeast genome organization, we can then ask where exactly replication occurs in the 3D nuclear space along the S-phase.

#### 1. A replication “wave” occurs across the nucleus

We start by investigating the localization of replication forks inside the nucleus. In particular, we compute the nuclear density of forks and monomers as a function of the 3D distance from the SPB for both interacting fork scenarios (Fig. 4, Fig. S11, Video S3,S4).

**FIG. 4:**
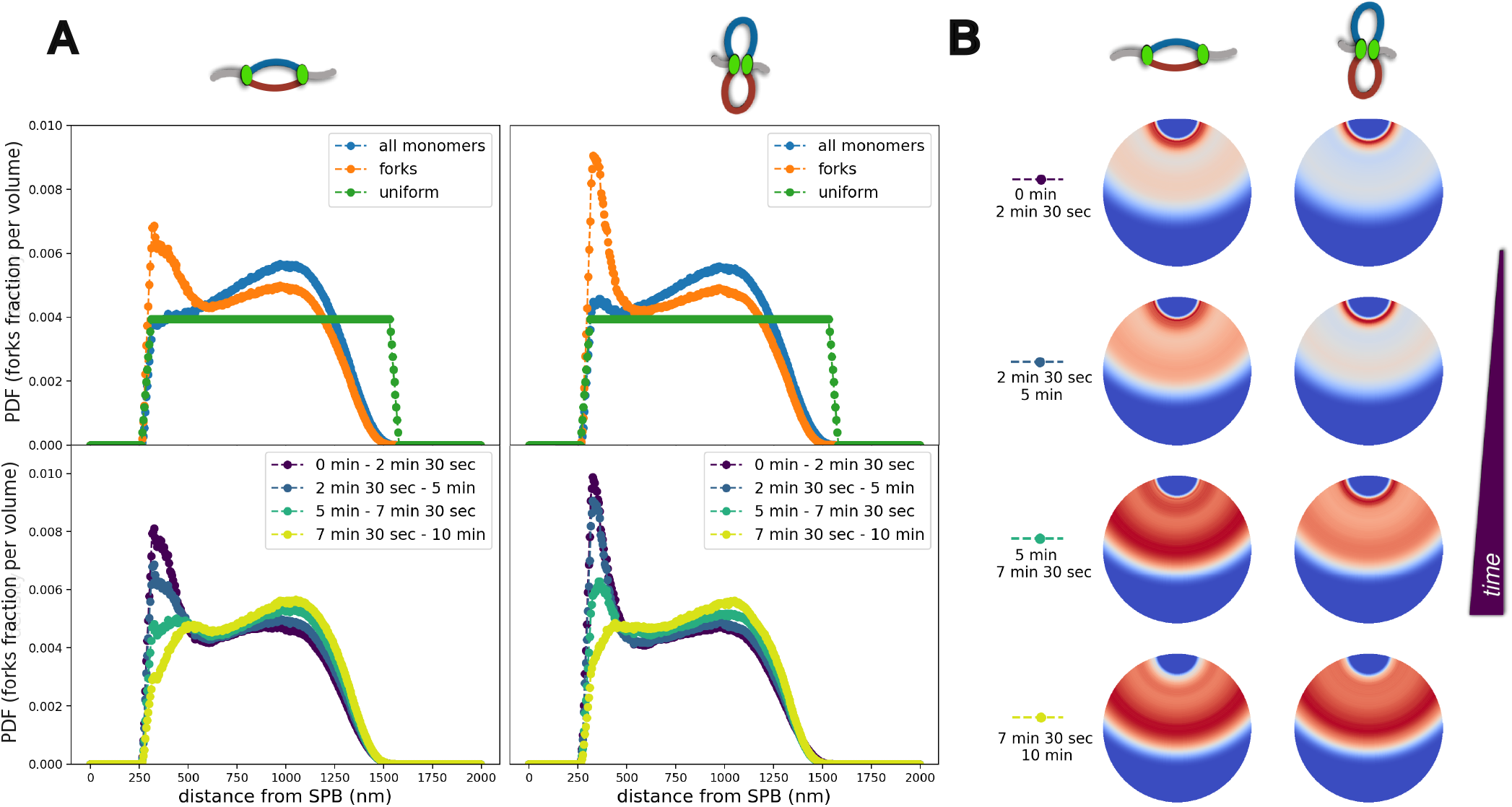
Forks spatial distribution. (A) Top: Probability Distribution Function (PDF) for a monomer to be at a distance *r* from the SPB: for all the monomers (blue) or only forks (orange), and compared with a uniform distribution (green). The computation includes the simulation frames between *t* = 2 min and 30 sec to *t* = 5 min in the non-interacting (Left) and interacting (Right) sister-forks scenarios. Bottom: Same as (A) for replication forks at different time intervals. (B) 2D graphical representation of the time evolution of forks densities plotted in the Bottom panels of (A). Note that the color scale used to plot the density profile of each time interval is normalized between 0 and 1.

We observe that the spatial distribution of all the monomers is heterogeneous inside the nucleus due to the Rabl organization, with an enrichment around the equatorial plane (Fig. 4A, Top, blue lines). Interestingly, this overall density evolves during S-phase, particularly in the interacting fork case with a slight enrichment close to the SPB in early-S that gradually vanishes at longer times (Fig. S11E), an effect that is not present for non-interacting sister forks (Fig. S11D,E). This demonstrates that replication not only impact locally the organization around origins (see above) but leads to a global rearrangement of chromatin within the nucleus.

When we focus on fork positioning, we observe a strong enrichment in early S around SPB (Fig.4A Top, orange curves). This is consistent with the higher abundance of early firing origins close to the centromeres. Interestingly, such an effect is stronger in the interacting forks scenario. Such enhanced localization around centromeres is gradually lost as replication progresses, with a relocation towards the equatorial plane (Fig. 4A Bottom and B and Fig. S11). Indeed, at larger times, centromeric regions are likely being replicated and forks tends to propagate more in the direction of telomeres. Moreover, new firing events of non-centromeric regions may further increase this effect.

Our results demonstrate the presence of a replication “wave” during S-phase in yeast, as already suggested by others based on MRT and Hi-C data [51]. More precisely, our simulations indicate that, on average, starting from an enrichment of replication forks at one pole (the SPB) in early S-phase, forks are redistributed in mid S-phase within the nucleus with a similar distribution comparable to the overall average chromatin distribution (blue curves in Fig. 4A, Top). In late S-phase (Fig. S11A,B), when the number of forks is low (Fig. S5), we observe a preferential enrichment at larger distances from the SPB. Note that such a “wave” can also be observed at the level of an individual trajectory (Video S3,4) even if much more stochastic.

#### 2. Replication forks may form small foci independently of specific aggregating forces

In the previous section, we observe a complex and heterogeneous distribution of replication forks within the nucleus driven in part by the Rabl architecture. Here, we investigate whether this peculiar spatial organization can explain by itself the presence of replication foci (RFi), where forks colocalize, as observed experimentally [14].

##### a. Characterization of replication foci in silico

To define and detect RFi in each simulation snapshot, we consider the independent components of the corresponding adjacency matrix *A*_*i,j*_, where *A*_*i,j*_ = 1 if forks *i* and *j* are distant by less than a threshold distance *θ* (*A*_*i,j*_ = 0 otherwise) (See Materials and Methods). *θ* may represent the typical spatial resolution of any experiments that allow to “visualize” the replication fork. For example for fluorescence-based microscopy, we might expect *θ* to vary from 60 nm (super-resolution-like) to 200 nm (confocal-like resolution). For both scenarios of sister-forks association, we computed the distribution of RFi during early S-phase, considering all frames between 2.5 and 5 min (Fig. 5A,B) for several *θ* values.

**FIG. 5:**
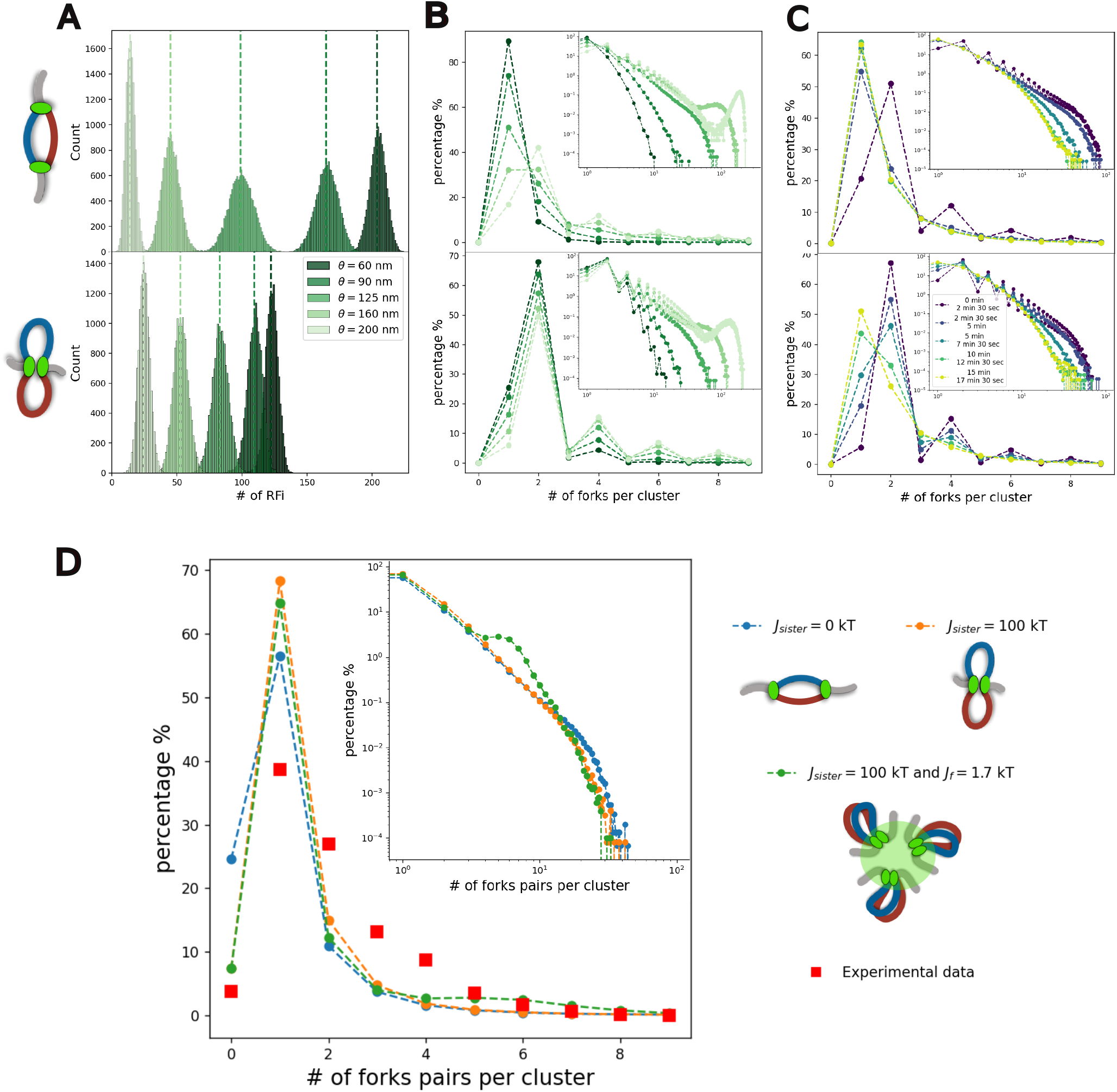
Detection of RFi within the nucleus during S-phase. (A) Distributions of the number of distinct RFi detected at a given resolution *θ* in early-S simulations (between *t* = 2 min and 30 sec and *t* = 3 min and 45 sec) for non-interacting (Top) and interacting (Bottom) sister-forks. The dashed lines correspond to the mean value of each corresponding distribution. (B) Percentage of clusters containing a given number of forks for simulations in early-S (between *t* = 2 min and 30 sec and *t* = 3 min and 45 sec) for non-interacting (Top) and interacting (Bottom) sister-forks and for varius resolutions *θ*. The inside panels show the distributions in logarithmic scale, including the probability for higher number of forks per cluster. (C) Percentage of clusters containing a given number of forks for different time windows in the non-interacting (Top) and interacting (Bottom) sister-fork cases. The inside panels show the distributions in logarithmic scale. (D) Left: percentage of clusters containing a given number of forks pairs. Red squares indicate the experimental estimates by Saner *et al*. (see Material and Methods). Simulated data in early-S (between *t* = 2 min and 30 sec and *t* = 3 min and 45 sec) was analyzed to strictly obtain even number of forks per cluster (see Material and Methods). Right: schemes and color relative to the three models explored in panel D.

In the non-interacting scenario (Fig. 5A, Top), at high resolution (*θ* = 60 nm), most of the individual forks can be resolved. Consequently, the average total number of detected RFi in the system is approximately equal to the average total number of replicating forks at that corresponding moment of S-phase. In the interacting sister-fork case (Fig. 5A, Bottom), as expected, at *θ* = 60 nm, the total number of RFis now corresponds mostly to the number of replisomes. As *θ* increases and the resolution is lowered, additional links are incorporated to the adjacency matrix *A*_*i,j*_, leading to a progressive shift towards lower values for the distribution of the numbers of RFi (Fig. 5A). Interestingly, for low *θ* values (≤ 160 nm), the interacting scenario always results in fewer detected RFi, each containing more forks on average due to the forced colocalization of sister-forks. However, for low resolution (*θ* ≥ 160 nm), observed RFi in the non-interacting case are less numerous (Fig. 5A) and larger on average. This is likely due to the more homogeneous distribution of forks in the non-interacting scenario (Fig. 4).

To further characterize the population of clusters, we compute the probability distribution of RFi sizes (Fig. 5B) at different resolutions *θ*. In the non-interacting case and at lower resolutions (*θ <* 125 nm), most detected RFi consist of a single diffusing fork (Fig. 5 B,Top). Increasing *θ* leads to the detection of some larger clusters. In particular, the probability of detecting sister-forks within the same cluster is enhanced, thus biasing the distribution to even numbers of forks (Fig. 5 B, Bottom, lighter green points). In the interacting forks, such a bias to even number is already present at high resolution (Fig. 5 B Top, darker green points) and the presence of clusters containing 4, 6 or 8 forks is not rare. Interestingly, at very low resolution (*θ* ≥ 160 nm), in both scenarios, very large RFis may be observed (Fig.S14C), containing a large fraction of the active forks (secondary peaks in the inset panels Fig. 5 B).

Fixing *θ* = 125 nm to mirror the optical resolution in the experimental work by Saner *et al*. (see below), we analyze how the sizes of RFis change during S-phase. In the non-interacting case, at very early S, the distribution shows again a bias in favor of even values (Fig. 5C Top panels, dark violet points) due to the small average replicon size and the subsequent proximity of sister-forks that are detected colocalized at such a resolution. The high density of forks at centromeric regions (as showed in Fig. 3C,D) increases also the probability of observing clusters (over 40% with 4 forks or more). This is rapidly lost as replication proceeds leading to detect a high percentage of single-forks (*>* 60%), and the gradual diminution of large RFis (Fig.5C top panels). In presence of sister-fork interactions, the bias towards even cluster sizes persists until 5 - 7 minutes (Fig.5C Bottom panels) after the beginning of S-phase. After, termination events where nearby replication bubbles merge lead to the loss of sister-fork interactions and thus to an increasing number of RFi with single fork (Fig.5C bottom panels, yellow and green points).

In summary, our *in silico* predictions indicate that random collisions of diffusing forks or sister-fork pairs may indeed lead to the significant detection of RFi containing several forks or replisomes. Clusters of large sizes are more abundantly detected in early-S when forks spatial distribution is enriched around the SPB (Fig. 3C,D).

##### b. Comparison with experiment

We test whether the predicted, resolution-dependent colocalization of multiple freely-diffusing forks is consistent with experimental observations made by Saner *et al*. [14]. Using super-resolution microscopy (SIM technology) on GFP-tagged PCNA in early S, a key component of replisome, the authors identified RFi at different time points along S-phase. For one time point where the total number of RFi is maximal, they inferred the number of sister-fork pairs inside a given RFi based on its overall fluorescence intensity (red squares in Fig. 5D). Note that such an inference was based on a *in silico* estimation of the total number of forks [73] and on assuming that sister-forks always remain associated (see Material and Methods). While in both of our scenarios, odd numbers of forks can be detected (Fig. 5C), we mirror the data analysis performed by Saner *et al*. by strictly casting the numbers of forks within RFi to even numbers (see Material and Methods).

In Fig. 5D, we estimate the distributions of the number of fork pairs inside our detected RFis (for *θ* = 125 nm, the typical spatial resolution of 3D-SIM [74]) at a time (2-4 minutes after the start of S-phase simulations) where the total number of replicating forks *in silico* is maximal (Fig. S5) and comparable to the estimation made by Saner *et al*. at the time of their measurements (~ 240 excluding rDNA-related forks). Interestingly, in our simulations, under these conditions, the average number of detected RFis is ~ 80 (respectively ~ 100) in the interacting sister-fork scenario (resp. non-interacting scenario), very close to the experimental observation (~ 75). Both scenarios of sister-forks association exhibit similar trends with a peak at 1 pair per cluster, as observed experimentally, and follow by a rapid decrease for higher number of pairs, failing to capture the significant probability of having RFi of intermediate sizes (2 - 4 pairs) inferred by Saner *et al*.

We thus wonder if such an underestimation of detected clusters of intermediate sizes *in silico* may be explained by the presence *in vivo* of aggregating forces that might stabilize larger clusters. Therefore, in addition to sister-forks interactions, we included non-specific pair-wise contact interactions between all the forks (see Material and Methods, Fig. S12,13). At the Hi-C level, we predict an enhancement of contacts between early origins (Fig. S12) as the strength of non-specific interactions *J*_*f*_ increases, that becomes significant for *J*_*f*_ ≥ 1.75 kT. At the level of the distribution of RFi sizes (Fig. S13), we observe only an enrichment of large clusters (5 - 15 pairs) for *J*_*f*_ = 1.75 kT, intermediate sizes (3 - 4 pairs) becoming more frequent for stronger *J*_*f*_.

Experimental Hi-C data in early S (Fig. S11D) do not exhibit significant contacts between early origins, suggesting that *J*_*f*_ *<* 1.75 kT. However at such interaction strength, the predicted frequency of intermediate RFi sizes are not compatible with microscopy data (Fig. 5D, S13).

In summary, the colocalization of replicating forks into RFi predicted by our modeling, either just by random collisions or mediated by aggregating non-specific forces, are not compatible with the super-resolution microscopy data of Saner *et al*. (see Discussion and Fig.S14).

### D. Replication impacts chromatin mobility

Finally, to complete the spatio-temporal description of replicating chromosomes, we investigate the impact of replication on chromatin mobility.

#### 1. Overall slowing down of chromatin diffusion due to catenations between sister chromatids

From simulated trajectories, we first estimate the average mean squared displacement of genomic regions (MSD, see Material and Methods) in G1 and along S-phase, for both sister-fork scenarios (Fig. 6 A, Fig. S15). Before the onset of replication, we observe the expected dynamics for a Rabl-like organization (Fig. 6 A, Fig. S13A) with an experimentally observed, Rouse-like, dynamics [75] characterized by a diffusion exponent 0.5 at short and intermediate time-scales [76, 60], which transitions to a more confined dynamics at larger times due to the various geometrical constraints imposed by the Rabl organization. As S-phase progresses, we observe an overall significant reduction of the diffusion constant (Fig. 6A, Fig. S15C) for both scenarios (Fig. S17). Interestingly, such a decrease is maximal between ~ 15 - 25 minutes when more than 65% of chromatin has been replicated and the number of active forks is rapidly decreasing (Fig. S5). At larger time, when most of the DNA has been replicated and with only few remaining replicating forks in the system, the reduction in diffusion constant is still present.

**FIG. 6:**
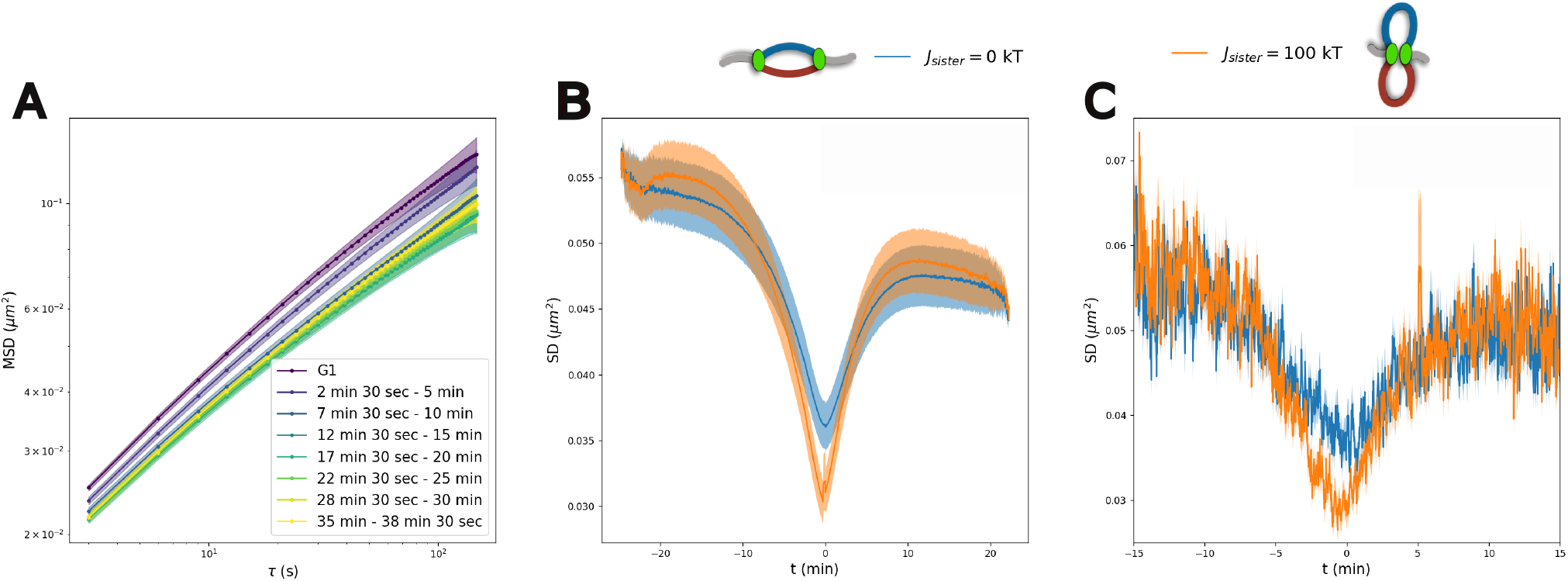
Chromosome dynamics during S-phase. (A) Mean Squared Displacement (*MSD*) for various time windows in S-phase in the case of non-interacting sister forks (*J*_*sister*_ = 0 kT). Curves correspond to an average across all the monomers and all the simulated trajectories. (B) Average Square Displacement *SD*(*t, τ* = 15 *sec*) for both scenarios of sister-forks association as a function of the time. *t* = 0 corresponds to the time of replication of the monomer. Curves correspond to an average across all the monomers and all the simulated trajectories. (C) *SD*(*t, τ* = 15 *sec*) for a single locus (Chromosome 2, 470 kb from left telomeres) averaged over all simulated trajectories.

We thus wonder about the physical origin of this global reduction in mobility. As it is strong even at times when very few, if any, forks are present, it cannot be attributed to the presence of actively replicating forks. Two possibilities remain: (i) an increase in the volumic density as, in our model, it is doubling during replication; (ii) the emergence of catenated structures from the intertwining of sister chromatids (Fig. S15D) [19]. To test these hypotheses, we allow all the chromosomes to replicate except one (chromosome 8, Fig. S16B), thus still doubling the overall volumic density. The MSD of the unreplicated chromosome is as in G1 while the other chromosomes exhibit a perturbed dynamics. Conversely, we allow only chromosome 8 to replicate (Fig. S16C). Similarly, the MSD of the replicated chromosomes is strongly perturbed while the unreplicated chromosomes behave as in G1. This suggests that changes in the volumic density - at least under physiological conditions - is not impacting chromatin mobility while the presence of catenated chains produced by the replication process slows down the mobility of individual loci [19].

#### 2. Fork passage transiently reduces dynamics

As mentioned above, the overall changes in *MSD* observed during S-phase are not caused by the number of active forks in the system or by the mode of sister-fork interaction. However, the non-trivial topologies at forks are expected to locally impact diffusivity [19] consistently with polymer theory of branching points [77]. We thus investigate if some dynamic signatures of fork passage may be observed in live-imaging data. For a given monomer, along a simulated trajectory, we compute the evolution of its Squared Displacement *SD*(*t, τ* = 15 *sec*) during *τ* = 15 seconds as a function of time *t* (see Material and Methods). We then compute the mean values of squared displacements around the replication time of this locus (Fig. 6B,C and Fig. S18). Averaging over all the loci, we observe significant decrease in diffusion at the moment of fork passage (*t* = 0) in both scenarios, with more pronounced reduction for the interacting case where forks are more constrained (Fig. 6B). Remarkably, our analysis suggest that, given enough statistics, such a behavior can be measured even by monitoring a single locus (Fig. 6C) in multiple cells or multiple loci in one single cell (Fig. S18B), thus opening the possibility to test this prediction experimentally by live-imaging.

## III. CONCLUSION AND DISCUSSION

In this study, we investigate the spatio-temporal organization of eukaryotic replication in the context of budding yeast nucleus. Our approach couples biophysical and computational modelling, with new *in vivo* HiC data in S-phase. Building on our previous work [19], we were able to contextualize our coarse-grained polymer model of replicating chromatin to the specific case of budding yeast, combining chromosomes’ large scale Rabl organization with an accurate description of the underlying 1D replication dynamics. As a result, our framework shows very high predicting power of chromosome folding during S-phase enabling us to quantitatively investigate the 3D organization of eukaryotic replication.

Throughout this study, we address the highly debated architecture of diverging sister-forks during S-phase that may - or may not - remain tether together [18, 17]. Remarkably, in both hypotheses of non-interacting and interacting sister forks, the formation of replication bubbles leads to the local (10s of kb) reorganization of chromosomes around origins of replication. In the former case, a mild contact enrichment around early replicating origins is predicted due to entropic effects [19]. In the latter scenario, distinct “fountain” or “jet” patterns emerge via an effective loop extrusion process. While previous studies have observed such a signature around origins in experimental Hi-C data in yeast [19] and in mammals [16], it was not demonstrated that these patterns are replication-dependent and do not emerge from other concomitant mechanisms.

Conducting new HiC experiments in a synchronized population of cells in early S-phase, we confirm the presence of fountains around early origins (Fig. 3A) and demonstrate that they are replication-dependent. Remarkably, the detection of such signal is strongly enhanced by our accurate classification of early origins if compared to the average signal computed using currently known ARS positions (Fig. S3, S10). More importantly, we show that fountains were fully retained even in early-S cells depleted of cohesins therefore excluding potential biases arising from the concomitant cohesin-mediated loop-extrusion activity and sister chromatid cohesion [55, 51, 53, 54]. However, the enrichment of contact observed *in vivo* data does not have the same magnitude as in the model prediction in the interacting scenario. We explore a number of realistic conditions - such as as radius of capture, heterogeneity in the cell population and mixed scenarios of associated and non-associated forks (Fig.S7, S8) - which may reduce the strength of the simulated fountain-like signal and make it closer to the experiments. All these analyses suggest that in order to simulate fountain patterns comparable to experimental data, a significant fraction of sister-forks *in vivo* may not be interacting. This may be due to the limited stability of such an interaction or to the existence of unknown limiting factors that restrict the number of possible sister-fork interactions. Therefore, having the possibility to experimentally perturb sister-forks colocalization would provide a direct mechanistic proof of fountain formation. According to our predictions, such a perturbation should lead to a transition from a loop-extruding pattern (fountains) to a milder entropically driven compaction (typical signal in the non-interacting case). Over the years, several studies proposed Ctf4 as a putative driver of sister-fork oligomerization [16, 14, 72]. In this study, we present the first Hi-C experiment on Ctf4-deficient cells in S-phase. We still observe fountain-like patterns (Fig. 3A), suggesting no significant change in the ability of sister-forks to interact, and thus challenging the role of Ctf4 in this process.

An interesting advantage of our framework is the ability to go beyond bulk properties and inspect large-scale structures. In particular, thanks of our minimal model of the Rabl-organization, we were able to investigate the coupling between the Replication Timing Program and the underlying nuclear architecture, providing the first *in silico* evidence of a redistribution of forks across the nucleus from one pole to the other (Fig. 4, Fig. S11, Video S3,4). Our framework suggests that the previously proposed concept of “replication wave” [51] cannot be easily visualized at the single cell level. Interestingly, rather than a gradual shift of forks density in time away from the SPB, we observe their redistribution from the regions around SPB in early-S phase to the whole accessible part of the nucleus, consistently with the average DNA density, in mid S-phase and finally to the equatorial plane in late S-phase (Fig. S11). Interestingly, we observe a transient impact of the replication on the overall local chromatin density in the interacting scenario (Fig.4 A) via an enrichment close to SPB in early-S, that may be interrogated experimentally in the future.

Given the specificities of Rabl organization and the abundance of forks close to the SPB in early S-phase, we question whether the passive coupling between nuclear architecture and Replication Timing program could recapitulate the experimentally-observed higher-order assembly of forks into Replication Foci by Saner *et al*. in yeast [14]. In this hypothesis, the detection of larger RFi is explained by the random collisions of diffusing forks (or sister-forks), physically constrained by the underlying 3D chromosome folding, as proposed in the mammalian context by more recent studies [26, 15]. However, our *in silico* estimation, regardless of the interacting scenario for sister-forks, cannot recapitulate the high probability of having RFi of intermediate sizes (containing 2 to 5 sister-forks pairs) observed in super-resolution experiments. Even when introducing non-specific fork-fork interactions, we could only stabilize larger clusters without improving our prediction at intermediate sizes (Fig. S14). This incompatibility might be partially explained by the challenging comparison between simulation and experimental outputs. Indeed, experimental results depend on an approximate estimation of the total number of forks at the time of measurements made by Saner *et al*. and assume that the poorer resolution on the z-axis does not introduce significant bias in cluster detection. However, we show that misevaluation of fork numbers or misidentification of small and large clusters can quantitatively affect the outcomes of the analysis (Fig. S14). Recent progresses in microscopy technologies which allow to reach resolutions comparable to our model [26] would enable for a more straightforward comparison in future studies. Discrepancies between simulations and experiments may also originate from missing mechanisms in the modeling framework like cohesin-mediated loop extrusion that may facilitate the clustering of nearby replisomes along chromosomes or the selective clustering of early origins mediated by the transcription factor Forkhead that has been shown to promote the firing of about 70-100 early origins and to create preferential 3D interactions between them [41].

To fully characterize the spatio-temporal impact of replication, our model was instrumental in exploring the dynamics of replicating chromosomes at a genome-wide level. We predict a general decrease in chromatin mobility progressively happening during S-phase due to the emerging catenation of the two SCs [19] rather than the presence or absence of forks or changes in volumic density (Fig. S16). It would be intriguing to test this hypothesis with live-cell imaging on synchronized cells during S-phase in absence of cohesins to prevent confounding effects from cohesin-dependent loop extrusion or cohesion. Moreover, such experiments would give insights on the possible role of Topoisomerase *in vivo* in releasing the intertwining between SCs. Locally, we quantitatively demonstrate that fork passage slows down dynamics and that this can be monitored experimentally if both the displacement and replication status of a given genomic locus are simultaneously recorded (Fig. 6B,C) as done previously [14, 13]. By aligning measured trajectories to the time of replication, our simulations suggest that given enough cells (~ 100s) a decrease in dynamics may be observed (Fig. S18A) at the time of replication and that the magnitude of such decrease is informative on the structure of the replisome.

Naturally, our minimal coarse-grained model cannot capture the full complexity of the system. In particular, similar to other modeling studies [58, 59, 60, 61, 62], we aimed to recover the core features of Rabl organization rather than a full accurate description, for example, of telomeric and centromeric regions. This limitation is illustrated by our poorer prediction for the last points of the *P*(*s*) curves (Fig. S2) or the predicted faster dynamics for telomeres (Fig. S15B). This is likely distinct from the complex behavior observed *in vivo*, where telomeres can be found both at the periphery and at the center of the nucleus and can aggregate through binding of the Sir complex and have a slower mobility [35, 32, 76]. In order to simplify our implementation, we have also chosen an intermediate nucleus radius (1 *µ*m) throughout replication, thus neglecting the experimentally observed gradual increase of size during S-phase [78]. Overall, we expect these limitations to have minimal impact on our results, mostly focused on early-S replication when less than 20% of the genome has been replicated.

Furthermore, to limit possible ambiguity and focus on replication-driven rearrangement of chromatin folding, we did not integrate possible interplays with SMCs. For this very reason, we performed HiC experiment on Scc1-depleted cells to isolate replication-dependent effects on 3D structure from the drastic rearrangements mediated by the cohesin complex in S-phase [55, 51, 53, 54, 45, 46]. Given the now well-established relevance of SMCs activity for genome maintenance and functioning all across the tree of life and cell cycle [3, 79, 80], it is crucial to investigate how it interferes with the core process of DNA replication. In the context of eukaryotes, despite recent advancement in the field [53, 43, 16, 81, 49, 82], it remains highly elusive to decipher how replication forks and different pools of cohesins, such as loop-extruding or cohesive, coexist in S-phase. In this perspective, our model could be instrumental to explore the possible mechanisms for cohesion establishment upon fork passage [44, 45, 47, 48]. Similarly, equipped with a loop-extrusion dynamics [83, 84] alongside replication [27], our model could provide mechanistic insights on the 3D effect of cohesin stalling [53], bypassing or detaching upon forks encounters.

Overall, our study suggests a significant impact of the replication process on the genome organization through an loop extrusion-like process mediated by sister-fork interactions and possibly through the formation of replication foci and a mild local and global reorganization of chromatin density. In yeast, where replication bubbles remain small, such an impact is transient and is not predicted to be persistent during G2/M. However, in species with larger genomes and replicon sizes [16], that may relax more slowly, we might expect a possible structural memory of the S-phase throughout G2.

What could be the biological functions of sister-fork interactions or replication foci on DNA replication itself? This question is still elusive as perturbing experimentally such processes and observing the consequences are still challenging. Integrating in our modeling framework a feedback of the 3D chromatin organization on the 1D replication dynamics via, for example, the 3D dependency of firing rates of origins or fork progressions would allow to test different mechanistic hypotheses [25, 85].

## IV. MATERIAL AND METHODS

### A. Experimental data

### 1. Haploid S. cerevisiae strains

Genotypes of *Saccharomyces cerevisiae*, (W303 RAD5+ background) strains are listed in Table S2. The *ctf4::KanMX* mutation has been obtained by transformation of a PCR fragment amplified from a KanMX cassette-containing plasmid using primers 5’-GTTTCCTGAATACGCCAACATATGGGAACATATAGATTAAATTAATAAGAAAGCTTGGGAcggatccccgggttaattaa-3’ and 5’-TTGAACGATGATTTGAACAAATGAACAGGTATCAAATAATTGTCTCTTGCGTATATATATgcgcgttggccgattcatta-3’ [86]. The construct *his3::pADH1-OsTIR1-9Myc::HIS3* for OsTir1 E3-ubiquitin ligase expression, as well as the Scc1-V5-AID and Cdc45-FlagX5-AID constructs have been described previously [50, 87].

### 2. Culture media and growth conditions

Cells arrested in G1 with alpha-factor in YPD medium (1% yeast extract, 2% peptone, 2% glucose) at 30°C were washed 3 times with 200 mL of pre-warmed YPD and released in S-phase at 30°C for 20 minutes prior to crosslinking. For Scc1-AID depletion and Cdc45-AID depletion, 2 mM IAA was added 2 hours prior to G1 arrest, and in all following wash and culture media.

### 3. Hi-C

Hi-C was conducted as described in [88] with minor modifications. Briefly, ~ 1.5 *·* 10^9^ haploid cells were fixed with 3% formaldehyde (Sigma-Aldrich, cat. F8775) for 30 minutes at RT at with orbital agitation at 120 rpm. Formaldehyde was quenched with 330 mM glycine for 20 minutes at RT at 120 rpm. Cells were washed twice with cold water at 3, 000 g for 10 minutes. Pellets were split in two tubes and frozen at −80 °C. Cell pellets were thawed in ice, 7.5 *·* 10^8^ were transferred to Precellys VK05 tube, and lysed for 3 *×* 30 s at 6, 800 rpm. Between 2 and 5 *·* 10^7^ cells were processed for Hi-C using the Arima Hi-C+ kit (Arima Genomics, cat. A410079) following manufacturers’ instructions. The Arima Hi-C+ kit employs a dual restriction digestion (DpnII and HinfI) yielding a median fragment length of 108 bp in S. cerevisiae. DNA was fragmented into 300 − 400 bp fragments using Covaris M220 sonicator. Preparation of the libraries for paired-end sequencing on an Illumina platform was performed using the Thermofisher Collibri ES DNA Library Prep Kit for Illumina Systems with UD indexes (cat. A38606024) following manufacturer’s instructions. The library was amplified in triplicate PCR reactions using oligonucleotides corresponding to the Illumina sequence adaptors (5’-AATGATACGGCGACCACCGAGATCTACAC-3’ and 5’-CAAGCAGAAGACGGCATACGAGAT-3’) and Phusion DNA polymerase (New England Biolabs, cat. M0531) for 11 cycles. PCR products were purified with AMPure XP beads (Beckman-Coulter, cat. A63881) and resuspended in pure H2O. The Hi-C library is quantified using the Qubit DNA high sensitivity kit (Thermo Scientific, cat. Q32851) on a Qubit 2 fluorometer (Thermo Scientific, cat. Q32866). Library quality control, paired-end sequencing (2 *×* 150 bp) on Illumina NovaSeq6000 or NovaSeq X Plus, and data QC were performed by Novogene UK. Libraries used are listed in Table S3.

### 4. Hi-C read alignment and data normalization

Paired-end reads were aligned and contact data filtered using the Hicstuff [89] pipeline function in “cutsite” mode with a quality threshold of 20. Briefly, pairs of reads are aligned iteratively and independently using Bowtie2 in its most sensitive mode to the S. cerevisiae reference genome R64-2-1, obtained from the https://www.yeastgenome.orgwebsite. Each uniquely mapped read was assigned to a restriction fragment. Contacts were filtered as described in [90] and PCR duplicates were discarded. Data were binned at 1 kb using Hicstuff rebin function and converted to a cooler format with Hicstuff convert.

### 5. Flow-cytometry

Approximately 10^7^ cells were collected by centrifugation, re-suspended in 70% ethanol and fixed at 4°C for at least 24h. Cells were pelleted, resuspended in 1 mL of 50 mM sodium citrate pH 7.0 and sonicated 10 seconds on a Bioruptor. After washing, cells were treated with 200 *µ*g of RNase A (Euromedex, cat.9707-C) at 37°C overnight. Cells were then washed and incubated for 30 minutes with 1 mL of 50 mM sodium citrate pH 7.0 with 16 *µ*g of propidium iodide (Fisher Scientific, 11425392). Flow cytometry profiles were obtained on a MACSQuant machine and analyzed using Flowing Software 2.5.1.

### 6. Protein extraction and western blotting

Protein extraction and western blotting were conducted as described in [82]. Protein extracts for western blot were prepared from 5.107 to 108 cells. Cells were lysed in cold NaOH buffer (1.85 N NaOH, 7.5% v/v beta-Mercaptoethanol) for 10 minutes in ice. Proteins were precipitated upon addition of trichloroacetic acid (15% final) for 10 minutes in ice. After centrifugation at 15, 000 g for 5 min, the pellets were resuspended in 100*µ*L of SB++ buffer (180 mM Tris-HCl pH 6.8, 6.7 M Urea, 4.2% SDS, 80 *µ* M EDTA, 1.5% v/v Beta-mercaptoethanol, 12.5 *mu*M Bromophenol blue). Proteins were denatured upon heating 5 minutes at 65°C. Pre-cleared extracts were resolved on 12% precast polyacrylamide gel (Bio-Rad, cat. 4561043) and blotted on a PVDF membrane (GE Healthcare, cat. 10600023). Membranes were probed with mouse anti-AID antibody (clone 1E4, CliniSciences, M214-3) diluted at 1 : 1000for Cdc45-Flag-AID, an anti-GAPDH antibody (Invitrogen, MA515738) diluted at 1 : 10000, or an anti-V5/Pk1 monoclonal antibody (Fisher Scientific, cat. R960-25) diluted at 1 : 10000 for Scc1-V5-AID. Primary antibodies were revealed with an HRP-conjugated anti-mouse IgG antibody diluted at 1 : 10000 (Abcam, ab6789) using Immobilon Forte western HRP substrate (Merck, WBLUF0100) and a Chemidoc MP Imaging system (BioRad).

### B. Polymer model and simulations

#### 1. Null model

Each chromosome is modeled as a self-avoiding walk on a fcc lattice composed of *N* beads and dynamically evolving via a Kinetic Monte Carlo (KMC) algorithm similar to other studies [91, 92, 93, 94, 95, 19]. In addition to excluded volume, monomers are subjected to a standard potential to account for the chain bending rigidity:

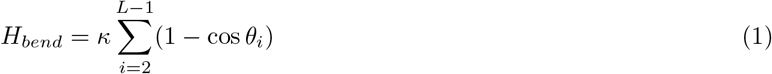

with *θ*_*i*_ being the angle between monomers *i* − 1, *i* and *i* + 1 and *κ* the bending modulus in kT units. To simulate chromatin, we consider standard values for the fiber diameter (*σ* = 20 nm) and compaction (50 bp/nm) [60], recovering an approximate bead size of 1000 bp and Kuhn length of 100 nm (*κ* = 3.217 kT). Time mapping between the simulation Monte Carlo Steps (MCS) and real time is done by computing the Mean Squared Displacement (MSD) of a monomer ⟨((*r*(*t* + *τ*) − *r*(*t*))^2^⟩ as a function of the time step *τ*. For this analysis, we used a single chromosome simulation of chromosome 4 of *Saccharomyces cerevisae* (polymer chain of 1531 monomers) in periodic boundary conditions and a 5% volumic fraction. Similarly to previous studies [93, 96, 97, 19] We then compared it to the experimentally-observed MSD_*exp*_(*τ*) [*µm*^2^] = ⟨((*r*(*t* + *τ*) − *r*(*t*))^2^ ⟩ ≈ 0.01 *· τ* ^1*/*2^ with *τ* in seconds, measured in *Saccharomyces cerevisae* [76], leading to 1 MCS = 0.075 msec.

#### 2. Full genome implementation

##### a. Initialization

Similar to previous analogous implementations [58, 60, 61, 62], we model the nuclear envelope by introducing a spherical barrier to the monomers in the system. In the lattice framework used here, this is accomplished by occupying all sites above a certain radius from the box center, now inaccessible to the polymer chains due to excluded volume interactions (Fig. S1A). Based on microscopy experiments [98, 35], we set the diameter of the sphere to 2*µ*m which corresponds to *≈* 70 lattice units given a bead size of 20 nm (Fig. S1B). We then introduce several distinct polymer chains into the system to model each individual chromosome. Since we aim to model haploid budding yeast genome, the number of chains is set to 17 where the 12th and 13th chains both model chromosome 12 (see below). Each chain is built using the HedgeHog algorithm [99, 93]: the chain is grown to the desired size (that could be different for each chromosome), starting from a simple backbone of length *L/*4 where *L* = 70 is the box size. Arbitrarily, we choose the backbone to be divided into two equally long arms, in random directions. The starting position of each backbone (corresponding to the first monomer of the chain) was placed at one of the 17 points inside the equatorial plane. These points are produced using the sunflower algorithm [100] to efficiently pack *n* points inside a circle (Fig. S1A, small panel). To randomize the process and avoid any biases, chromosomes are built in a random order, ensuring that the two random directions used to build the backbone of a new chain do not cross the ones already initialized. Note that such initial configuration does not aim to mimic any biological scenario (such as chromatids organization upon Mitotic exit). The choice of a such randomized V-shaped backbone, allows for simple and efficient construction of non-crossing chromosomes in the lattice, whose chain ends (telomeres) are randomly oriented. Given the strong forces that will be introduced in the system to recreate the Rabl configuration, this initial choice is likely to have a minimal impact. In fact, different initialization strategies in other studies [64] eventually converge to similar results once those forces are established.

##### b. Rabl organization

To mimic centromere attachment to the SPB, we introduce a spring-like potential between the centromere of each chromosome and position 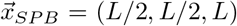 in XYZ coordinates (Fig. S1B). This forces the centromeres in the first times of the simulation to move towards the SPB (spring constant *k* = 100 kT). We complement our potential to allow the centromeres to freely diffuse in a spherical shell located between 250 nm and 325 nm from the SPB. The Hamiltonian describing the kinetic of centromeric monomers is:

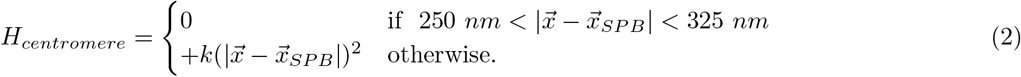

To simulate telomeres tethering to the nuclear envelope (NE), we implement an outward force acting on the terminal monomers of the chains, with a negative spring potential connecting them to the centers of the spheres 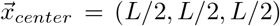 to push them towards the NE. Again, we allow the telomeres to freely diffuse within a distance of 50 nm from the spherical wall (Fig. S1B). Therefore, the Hamiltonian describing the kinetic of telomeric monomers is:

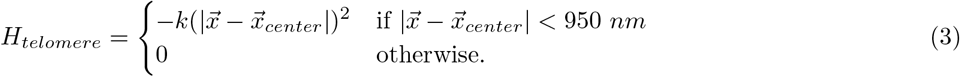

Since the nucleolus consists of a large physical barrier for non-rDNA, we introduce an additional impermeable wall positioned at 800 nm in the z-direction (Fig. S1B,C). Spherical confinement and the nucleolus wall lead to a final volumic fraction Φ = 3%, compatible with chromatin concentration standardly used for modeling yeast chromosomes [93].

##### c. Modelling chromosome 12

To simulate chromosome 12 and account for its role in the nucleolus assembly, we split chromosome 12 into two polymer chains anchored by one side to the nucleolus wall (Fig. S1C). The first chain models the first 460 kb of chromosome 12 where its rightmost monomer corresponds to a boundary of rDNA (Fig. S1C). The remaining portion of the chromosome, without centromere, is modeled with a second chain whose leftmost monomer constitutes the second boundary of rDNA, while the rightmost one, the second telomere. The two rDNA beads are pulled to the nucleolus wall with a spring potential only in the z-direction (downwards) enabling their localization at the nucleolus wall periphery:

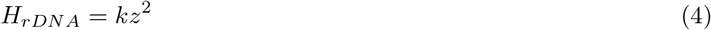

##### d. Comparison with in vivo HiC maps

Due to the brush architecture and centromere anchoring at the SPB, the two arms of a single chromosome tend to interact more if compared to an unconstrained polymer. This leads to a typical enrichment for the inter-arms signal (Fig. 1A,B) captured in both experiment and simulation. Moreover, we find an average Pearson correlation between experimental and simulated HiC maps at the single chromosome level (chromosome 4 in Fig. 1B) of 0.94.

We compute aggregate plots around centromeres (on diagonal plots, see below for technical details) normalized by the intra-chromosome *P*(*s*) (Fig. 1C). Our model correctly predicts the average shape of “flame” around centromeres in experimental maps even if we observe a too strong depletion of contacts between centromeres and chromosome arms, compared to *in vivo* maps (blue stripes in Fig. 1C).

We then quantify the enrichment of centromere-centromere (*trans*) contacts arising from the imposed colocalization of centromeres in the confining shell around SPB by computing off-diagonal aggregate plots. To compare the experiment and simulation, we normalize both by dividing by the respective *P*_*intra*_(*s* = 10 *kb*) (Fig. 1D). In the simulations, we estimate that the average centromere-centromere contact frequency is ~ 11% of the *P*_*intra*_(*s* = 10 *kb*) while the experiment exhibits a more moderate enrichment ~ 8%. In both cases, these quantitative differences are likely due to the strong modelling assumptions made. For instance, chromosomes are strictly bound by the strong potential to localize between 250 and 320 nm from the SPB. Relaxing these conditions should decrease the centromere-centromere contact frequency. Similarly, the strong potential brings telomeres and centromeres strictly to different parts of the nucleus (NE and SPB respectively). This polarity is enforced in all trajectories, bringing a very strong depletion of centromeres and chromosome arms (blue stripes in Fig. 1A). The segregation between centromeres and telomeres might be more heterogeneous *in vivo* and other biological processes (such as condensin activity) may hinder this effect.

#### 3. Self-replicating polymer

##### a. Generic model

We model explicitly replication in 3D using a specialized class of Self-replicating polymers to simulate each chromosome. On-lattice chain duplication in KMC simulations was physically characterized in details in our previous study [19]. While in this study, we include multiple chains and Rabl-organization, the local realization of replication is analogous since each polymer chain, while governed by a global underlying 1D replication dynamics (see below) duplicates “independently” in 3D. Briefly, a given unreplicated monomer can be used as an origin of replication, leading to the formation of a replication bubble. In particular, origin firing consists of introducing into the system a new monomer, connecting it to the two neighbors of an origin along the maternal chains. This process effectively creates two polymer branches, which locally simulate the two nascent SCs. Monomers with triple connectivity at the extremity of the replication bubbles are defined as replication forks with a given directionality based on their position with respect to the origin. With a constant speed *v* = 2.2 kb/min, every fork in the system can trigger further replication by introducing additional monomers at the fork positions and moving the triple connectivity. Replication bubbles merge once two forks of opposite directionalities converge along the chain. In such a configuration, the two converging forks, now neighbors along the chain, are always replicated in pairs, leading to the formation of a single bigger bubble. Similarly, when a fork is adjacent to a telomere, the bubble is opened by replicating both the fork and the chain end monomers.

##### b. 1D replication dynamics

We introduce implicit 1D replication dynamics by adapting the formalism of Arbona *et al*. [56, 57] to our KMC framework. Replication is modelled as a bimolecular reaction between a limited number of *N*_*f*_ firing factors and the assigned origins of replication and a firing trial move is introduced in our scheme. Under the assumption of a well-mixed system, at a given time, a set of *N*_*origin*_ origins is randomly selected (see below) and each origin of this set is fired with a probability:

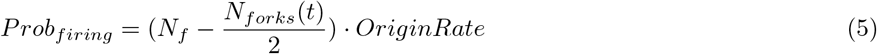

where (*N*_*f*_ − *N*_*forks*_(*t*)*/*2) represents the current number of available freely diffusing firing factors and *OriginRate* is a constant rate, equivalent of *k*_*on*_ in [56, 57]. To model the heterogeneous time patterns of eukaryotic replication, the set of *N*_*origins*_ is sampled from the “Initiation Probability Landscape Signal” (IPLS) computed in [57]. The IPLS is a signal that defines origin priming efficiency and was inferred in [57] through a machine learning approach to accurately reproduce experimental Mean Replication Timing and Replication Forks Directionality data [38]. Practically, the sampling of origins is performed by randomly picking a fixed number of monomers (*N*_*origins*_ = *L*_*chromosome*_*/*5 as inferred by Arbona *et al*. in [57]), the weighted probability of each monomer being given by the binned IPLS at the same resolution of our model (1 kb). Note that the drawing is done with replacement, meaning that each monomer can be selected multiple times to account that a certain coarse-grained genomic position may contain multiple (fine-scale) origins, which all have the same probability of firing (*OriginRate*). We then set *N*_*df*_ = 120 and *OriginRate* = 1 *·* 10^*−*9^*MC*^*−*1^ which are the KMC parameters equivalent to the ones inferred by Arbona *et al*. in [57]. Firing factors are rapidly introduced in the system once the S-phase starts according to the formula *N*_*df*_ (1 − 0.13 exp^*−t/τ*^) with *τ* = 1 min and 30 sec.

##### c. Interaction during S-phase

Co-localization of sister-forks is maintained by adding an elastic-like potential between the forks to the Hamiltonian of the system on which the Metropolis criterion is applied:

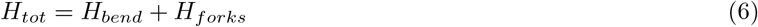

with

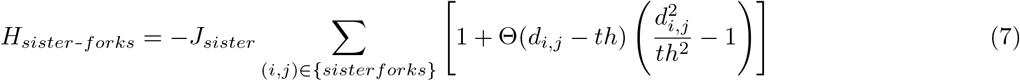

where the sum is performed over all the pairs of still active sister forks localized at monomers *i* and *j, d*_*i,j*_ is the 3D Euclidean distance between *i* and *j*, Θ(*x*) is the Heaviside step function and *th* = 50 nm is the distance below which the energy saturates to *J*_*sister*_ = 100 kT. Springs between forks are kept until one of the two interacting partners is lost due to the merging of replication bubbles or replication of the chain ends.

##### d. Non-specific forks interactions

We include contact interactions with strength *J*_*f*_ between all the replication forks in the system. Similarly to previous copolymer lattice models [93, 92, 96, 94], we include an additional term on the Hamiltonian of the system:

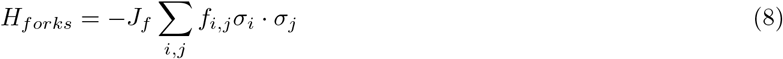

where *σ*_*i*_ = 1 if monomer *i* is a replication fork, otherwise *σ*_*i*_ = 0. *f*_*i,j*_ = 1 if monomers *i* and *j* are second nearest-neighbors otherwise *f*_*i,j*_ = 0.

The value of *J*_*f*_ = 1.7 kT was set by visually inspecting HiC maps (Fig. S12A) and off-diagonal aggregate plots (Fig. S12B) between early origins for increasing values of *J*_*f*_. In particular, we selected an energy value which lies in the transition between non-aggregated and aggregated states [92, 94].

### C. Data analysis

#### 1. Computation of HiC maps

The contact probability between any two monomers *i* and *j* is defined as the probability that the 3D Euclidean distance *d*_*i,j*_ between *i* and *j* is less than a fixed radius of contact *r*_*c*_. *r*_*c*_ is set to 80 nm in all the figures except Fig. S2A. The full genome contact matrix is then computed by ordering all the monomers according to their genomic sequence. The resulting matrix is loaded into a cooler file [101] at 1 kb resolution (monomer size) and then converted to a multi-resolution .*mcool* file to obtain the maps at lower resolution. In the presence of replicating DNA, we do not distinguish between SCs when populating the *in silico* HiC maps. As a result, similarly to the *in vivo* experiments, the contacts between two genomic positions *i* and *j* can be populated by different monomer pairs according to the current replication status of the two. Unless stated otherwise, contact maps (either *in silico* or *in vivo*) are always normalized using the function *balance*.*iterative correction* from the cooltools library (v 0.5.1) [101]. Expected contacts in *cis* (*P*(*s*) curves) and in *trans* are computed using the *expected* functions of cooltools [101]. Pileup plots are computed using the “pileup” function in coolpup library [102]. Specifics on the number of independent simulations and frames used for each plot are summarized in Table S1 in the Supplementary Materials.

To define early and late replicating origins, we used the IPLS from [57] (Fig S3) that code for the frequency of firing in our framework. We use the Scipy function *find peak* [103] to extract the peaks of IPLS. We define the 15% highest peaks (signal *>* 5) as early origins of replication and all the remaining peaks whose signal is above 1 as late origins.

#### 2. Comparison between in silico and in vivo signals

In Fig. 1A,B we rescale simulated raw contact matrices by multiplying the signal by a factor 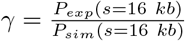. To compare simulated and experimental *P*(*s*) in Fig. S2B,C and Fig. S8, we multiplied simulated curves by a factor *α* which minimizes the Mean Squared Error (*MSE*) within a given genomic interval [*s*_*min*_, *s*_*max*_]:

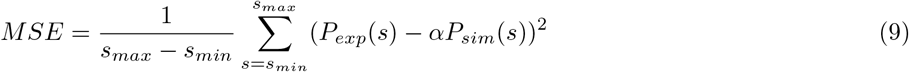

Practically we use *s*_*min*_ = 10 kb and *s*_*max*_ = 1 Mb when comparing *P*(*s*) averaged among different chromosomes (Fig. S2B and S8) while from *s*_*min*_ = 10 kb and *s*_*max*_ = 50 kb in the case of individual chromosomes (Fig. S2C).

#### 3. Spatial distribution of forks in the nucleus

In Fig. 4 and Fig. S11 we plot the Probability Distribution Function of forks as a function of the distance from the SPB. Summing over all trajectories, we count the number of forks at a distance between *r* and *r* + *dr* where *dr* = 5 nm and at a given time window. We then normalize by the total number of forks and the corresponding volume of the slice: *V* (*r* + *dr*) − *V* (*r*) where *V* (*r*) = (3*R* − *r*)*r*^2^*π/*3 is the spherical cap for a distance *r* from the SPB and *R* = 1000 nm is the radius of the sphere. We also compute the same observable for all the monomers to assess whether replication forks localization follows the average polymer organization.

#### 4. Analysis of RFi

##### a. Definition of RFi in silico

We define RFi *in silico* by constructing a graph based on the 3D positions of forks, 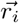, at each simulation frame. For a given threshold 3D distance *θ*, we define an association matrix *A*_*i,j*_, where *A*_*i,j*_ = 1 if 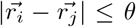 and *A*_*i,j*_ = 0 otherwise. An RFi is defined as an independent component of *A*_*i,j*_ (ie. a set of forks where each fork in the set is separated by a distance *> θ* from any forks outside the set). The total number of RFis corresponds to the number of graph components. To efficiently compute this quantity, we utilized the Python library *networkx* [104].

##### b. Comparison with experimental data

To compare our predictions to the experimental estimates of forks within

RFi by Saner *et al*. [14] (Fig. 5D), we adapt our cluster detection to mirror the experimental protocol. In the original paper [14], the authors analyze 3D microscopy images of replicating yeast nuclei where PCNA (forks subunit) has been labeled with GFP proteins. Each distinct bright spot detected is then classified as an individual RFi and the number of forks inside is computed on the basis of an *in silico* estimation of 302 forks in the nucleus at that specific replication stage. In practice, the relative intensity of each focus (signal of the focus divided by total signal) is multiplied by the total number and cast strictly to an even number. This assumption is motivated by early microscopy evidences that sister-forks are associated [13]. Experimental estimate of forks pairs within each RFi (red squares) was extracted from Figure 3E of the original paper [14]. In our *in silico* predictions, we roughly follow a similar approach by dividing the number of forks of each RFi by the total number of forks in the system in that specific frame to obtain a relative signal. For each RFi, we then multiply such value by the average number of forks between *t* = 2 min and 30 sec and *t* = 3 min 45 sec (229 forks). Finally, the resulting number was cast to even. In Fig. S14, we explore confounding factors that may impact the classification.

#### 5. Analysis of chromosome dynamics

In order to simplify the analysis, all the observables described below are computed used only one chromatid for each chromosome. In particular, once a given monomer is replicated, each measurement is computed only for its copy belonging to the Sister Chromatid 1.

##### a. Mean Squared Displacement

In Fig. 6 and S15C,17, we compute the Mean Squared Displacement with the formula:

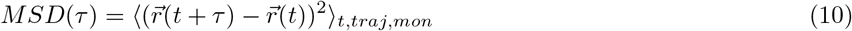

In Fig. S15B, we average over the monomer (*mon*) in the centromeric or telomeric regions, defined as being at a genomic distance less than 20 kb of a centromere or telomere respectively. Similarly, in Fig. S16 the MSD is computed restricting the average to monomers belonging to individual chromosomes. In Fig. S16B, we impaired the replication of chromosome 8 by manually deleting all the potential origins before the onset of replication. Conversely, “instantaneous” replication was achieved by setting a homogeneous high firing rate for all origins and high replication speed (Fig. S16C) for all chromosomes or exclusively for chromosome 8 (and hindering replication for all the others, Fig. S16D).

##### b. Time-rescaled Squared Displacement

The average Squared Displacement at 15 seconds in Fig. 6B, was computed with the formula:

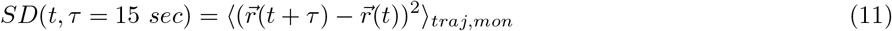

To rescale time, we recorded each individual *SD* trajectory along with its replication status (see Fig. S18A for an example of a single monomer in a single simulation) and aligned all trajectories at the moment of replication (*t* = 0 min) and eventually compute average trajectory (Fig. S18A).

## ACKNOWLEDGEMENT

We are grateful to Maxime Tortora, Geneviéve Fourel, Benjamin Audit, Maria Barbi, Amith Zafal Abdulla and members of the Jost lab for fruitful discussions. We acknowledge Agence Nationale de la Recherche (Grants No. ANR-18-CE45-0022 for D.J., ANR-21-CE45-0011 for C.V. and D.J. and ANR-23-CE12-0014 for A.P. and D.J., ANR-23-CE45-0033 for J-M.A.), the European Research Council (ERC) under the European Union’s Horizon 2020 for A.P. (ERC grant agreement 851006) and ENS de Lyon (D.D.) for funding. We thank Centre Blaise Pascal de simulation et modélisation numérique of the ENS de Lyon for computing resources.

## DATA AND CODE AVAILABILITY

Simulation code is available at https://github.com/physical-biology-of-chromatin/LatticePoly/tree/ master on the branch named *Y east*_*f*_ *ull*_*g*_*enome*. Hi-C data are available at the GEO repository numbers indicated in Table S3 *(unpublished data will be uploaded after the manuscript is accepted)*.

## AUTHOR CONTRIBUTIONS

D.D., J.-M.A., C.V. and D.J. designed and led the project. D.D. developed the polymer model and data analysis pipelines. D.D. performed simulations and data analysis. J.-M.A. inferred IPLS for the replication dynamics. V.P. and A.P. performed experiments. D.D., A.P., J.-M.A., C.V. and D.J. analyzed and interpreted the results. D.D. and D.J. wrote the manuscript with inputs from all the other authors.

