## supplementary materials for "Genome-wide modeling of DNA replication in space and time confirms the emergence of replication specific patterns in vivo in eukaryotes"

This document contains:

- Supplementary Figures S1 to S19
- Supplementary Tables S1 to S3
- Captions of Supplementary Videos S1 to S4

### Supplementary figures

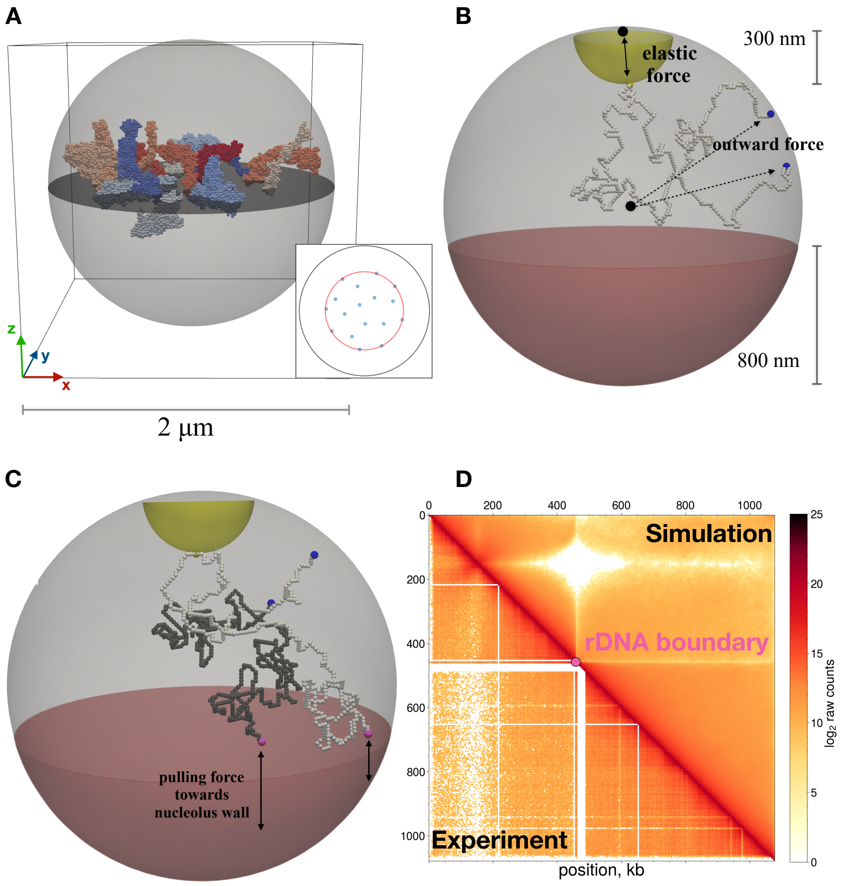


Figure S1: **On lattice implementation of the yeast Rabl-organization** . (A) Example of an initialized system on the lattice. The gray area indicates the volume accessible to monomers. (Inset) The starting monomer of each chromosome for the hedgehog algorithm is positioned in one of the blue points within the red circle located at the equatorial plane. (B) Example of a chromosome in a Rabl configuration. The yellow monomer indicates the centromere confined in the yellow shell to mimic SPB attachment. Blue monomers correspond to the telomeres pushed towards the NE. The red area indicates the nucleolus, whose volume is assumed to be inaccessible to monomers. (C) Example of a configuration of Chromosome 12 on the lattice. The light gray polymer chain models the first 460 kb of DNA. The rDNA boundaries (indicated by pink monomers) are pulled in the $-z$ direction. The darker polymer chain models the rest of Chromosome 12 without centromere. (D) Comparison between experimental and simulated contact maps for chromosome 12.

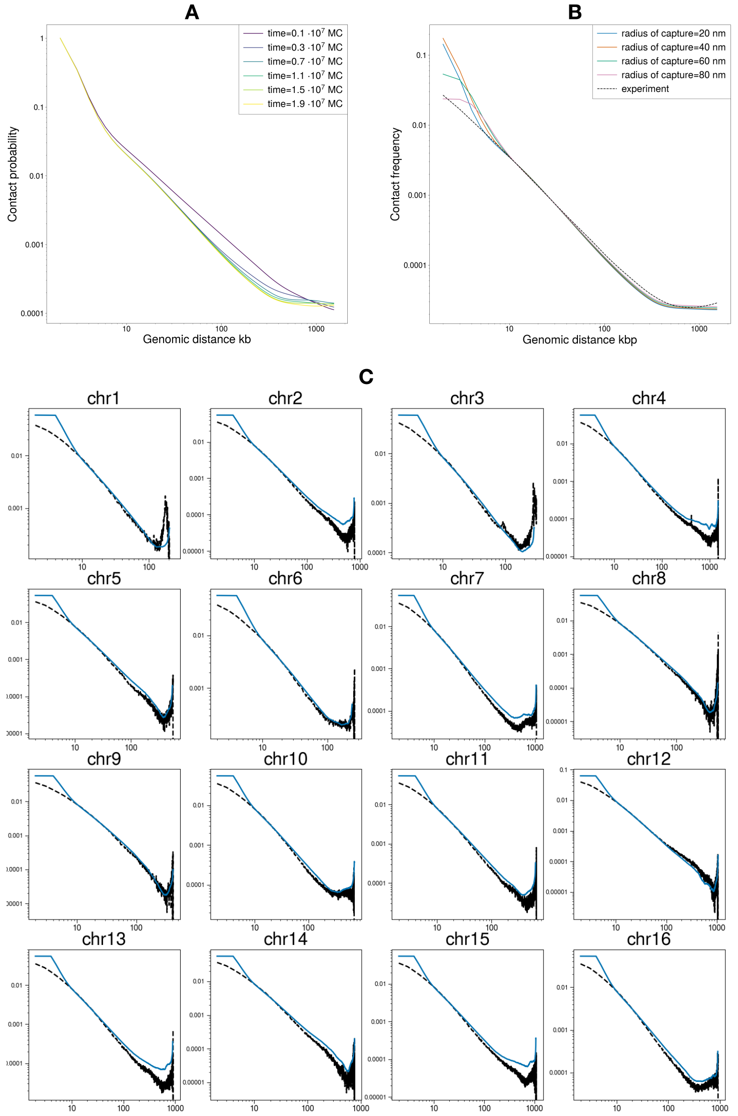


Figure S2: **Comparison with *in vivo P(s)* curves.**(A) Since the initial configuration in Fig. S1A does not aim to capture any biological phenomenon, we need to establish a relaxation time required to reach a G1-like 3D chromosome organization. While centromere, rDNA and telomere monomers reach their constrained locations in a few MC steps, each chromosome has to equilibrate its internal distances before making any measurements. *(Continue in next page)*.

Figure S2: *(Continued)* For this reason, we investigate intra chromosome $P(s)$ by computing contact maps at various time steps of the relaxation process (radius of contact of 40 nm) establishing a relaxation time of $10^7$ MCS. The $P(s)$ signal was smoothed and averaged between all chromosomes using *cooltools* functions. In this case, no normalization is applied (raw contacts). (B) $P(s)$ curves for 4 different radii of capture (20, 40, 60,80 nm) after averaging over all the chromosomes. Simulated curves are rescaled by multiplying by a constant factor $\alpha$ computed as described in the Materials and Methods ($s_{min} = 10$ kb and $s_{max} = 1$ Mb). Interestingly, independently of the chosen radius of capture, our *in silico* curves well recapitulate the experimental scaling of intra-chromosome $P(s)$ from G1 HiC experiment. However, our model does not capture very well the contact frequency at a large scale. In particular, in the smoothed experimental data (Fig. S2B black dashed curve), we can observe a small increase in contact at the end of the curve due to telomere-telomere contacts.(C) Simulated (blue, $r_c = 80$ nm) and experimental (black) $P(s)$ curves for all 16 chromosomes. Simulated curves were rescaled multiplying by a constant factor $\alpha$ where $s_{min} = 10$ kb and $s_{max} = 50$ kb. For every chromosome, the simulations capture the increase of contact frequency at the end of the curve due to telomere clustering. However, in our simulations such an increase is much steeper (see chr1 or chr3 for a more visually clear example), likely due to more rigid constraints by our modelling. In particular, telomere monomers are always strictly tethered to the NE, while they may also be observed in the interior [1]. We speculate that the lack of such heterogeneity at a larger scale in our model might result in the flat $P(s)$ at large scales once averaging all the chromosomes and smoothing (Fig. S2B).

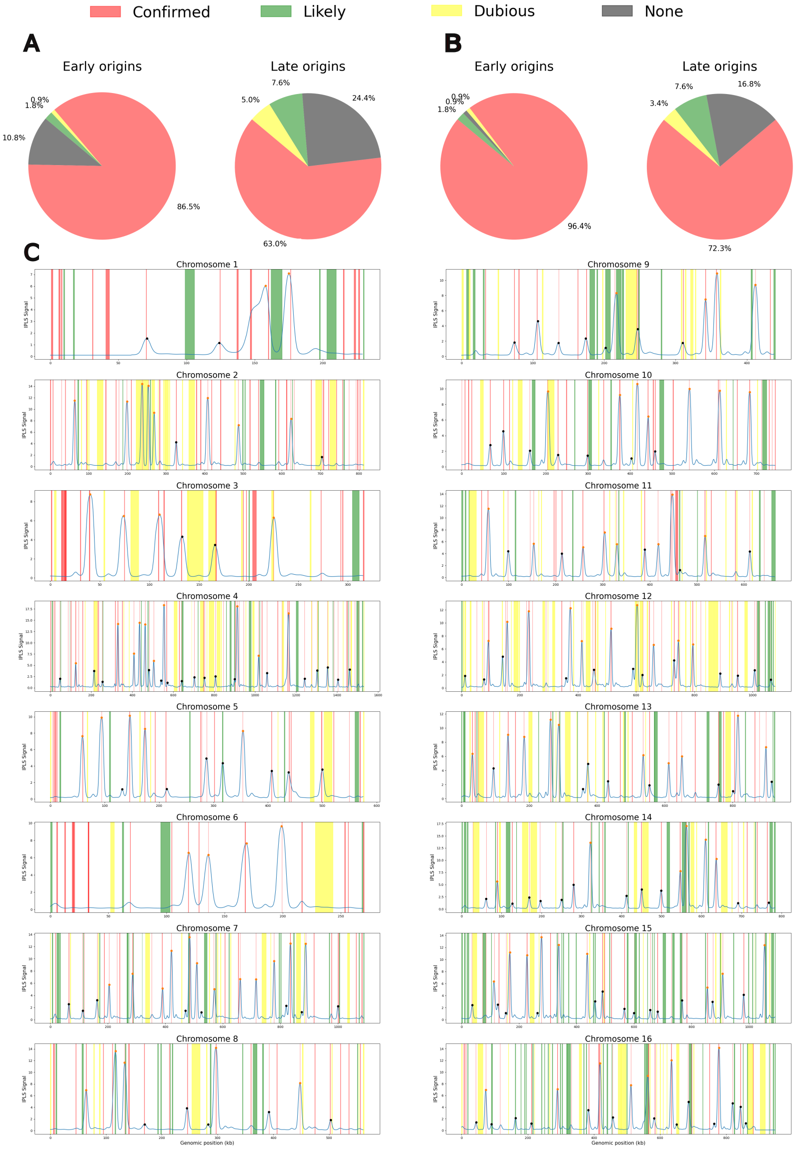


Figure S3: *(Caption in the next page.)*

Figure S3: *(Continued)* **Comparison with Oridb ARS.** (A,B) Percentage of origins detected through the $IPLS$ (See Materials and Methods for more details) which coincide to known ARS in budding yeast from the Oridb database [2]. According to the nomenclature in Oridb, each origin corresponds to either a "Confirmed" (red), "Likely" (green) or "Dubious" ARS or "None" if no match was found. When multiple ARS match we use the best classification available. In (A), we assign origins to a class when the position inferred from the $IPLS$ is at most 1 kb distance to an ARS, considering the start and end in the sequence from Oridb. On the right we used the origin in this study referred as "Early" (see orange circles in panel B) while on the left, "Late" origins exclusively (see black circles in panel B). (B) Same as (A) relaxing the classification with a genomic distance between our origins and ARS of 2 kb. (C) $IPLS$ profiles and inferred origin positions (peaks) for all chromosomes. The positioning of "Confirmed" (red), "Likely" (green) or "Dubious" ARS from Oridb is also shown. In conclusion, our method is very accurate in predicting origins positioning, since a high percentage was found to correspond to "Confirmed" ARS. Interestingly, several confirmed ARS do not match with a peak of the $IPLS$ signal, which is computed solely using *in vivo* MRT and RFD data [3].

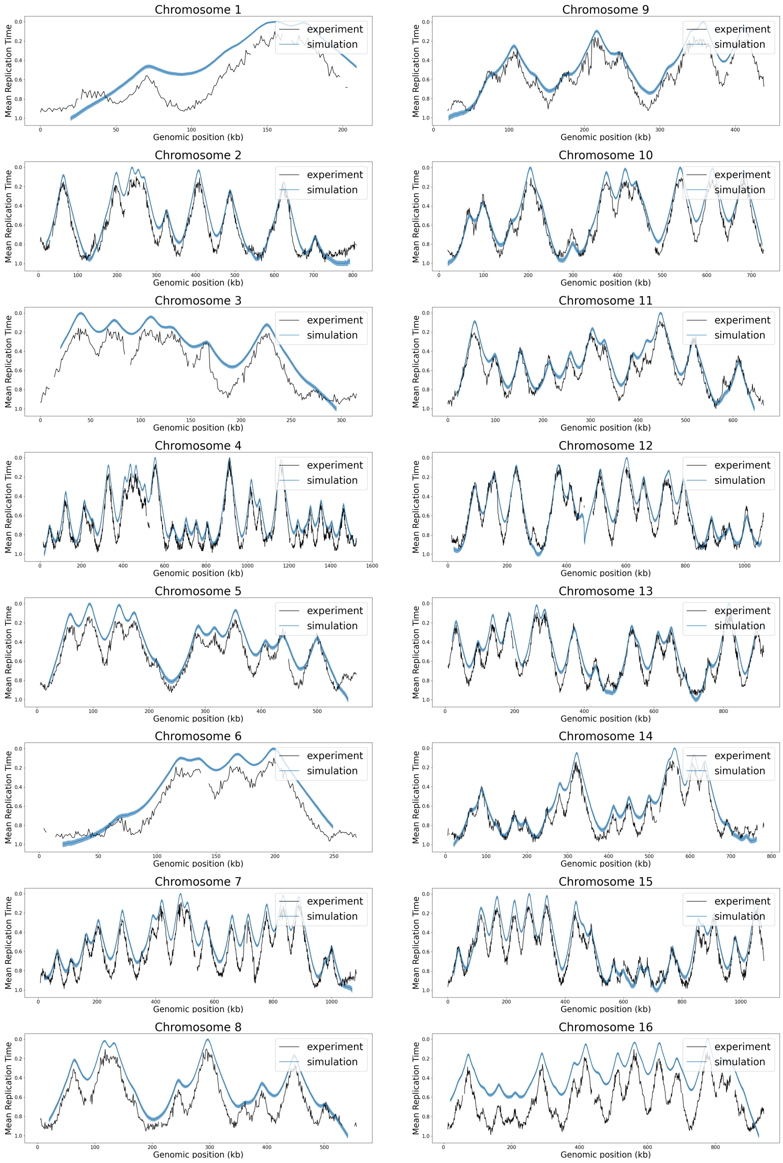


Figure S4: **1D Replication dynamics**. Comparison between the experimental data [4] (black lines) and simulations (blue lines) for all the chromosomes. We computed the simulated MRT by rescaling the average replication time of each monomer between 0 and 1. We excluded in the computation the last 20 kb of DNA which are not well captured by the model. Slow replication of telomeric regions can strongly affect the signal once simulated MRT is rescaled as it can still be observed for chromosome 16 where we speculate that the IPLS fails to capture origin firing close to the telomeres.

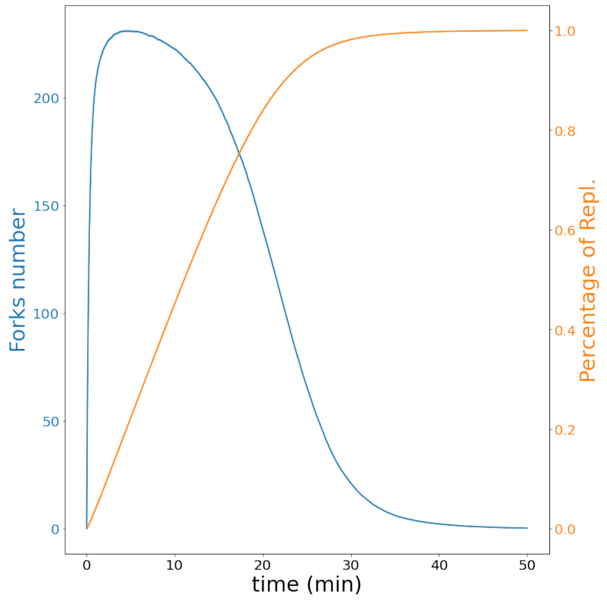


Figure S5: **Average number of forks and percentage of replicated chromosomes over time predicted by the model over the whole genome**.

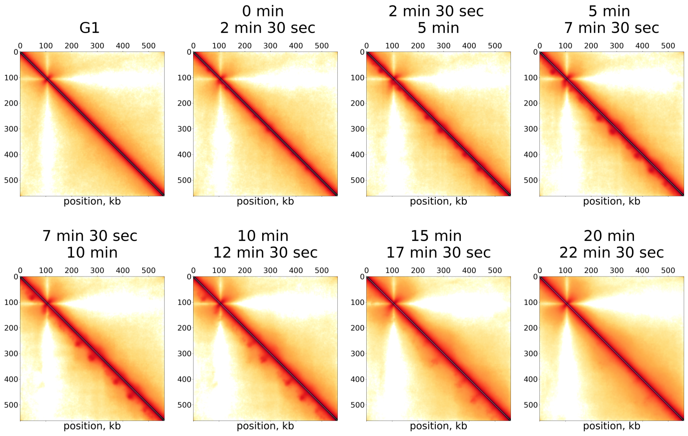


Figure S6: Example of normalized HiC maps (chromosome 8) at different time steps in the S-phase for interacting (bottom triangle) and non-interacting (top triangle) sister-forks.

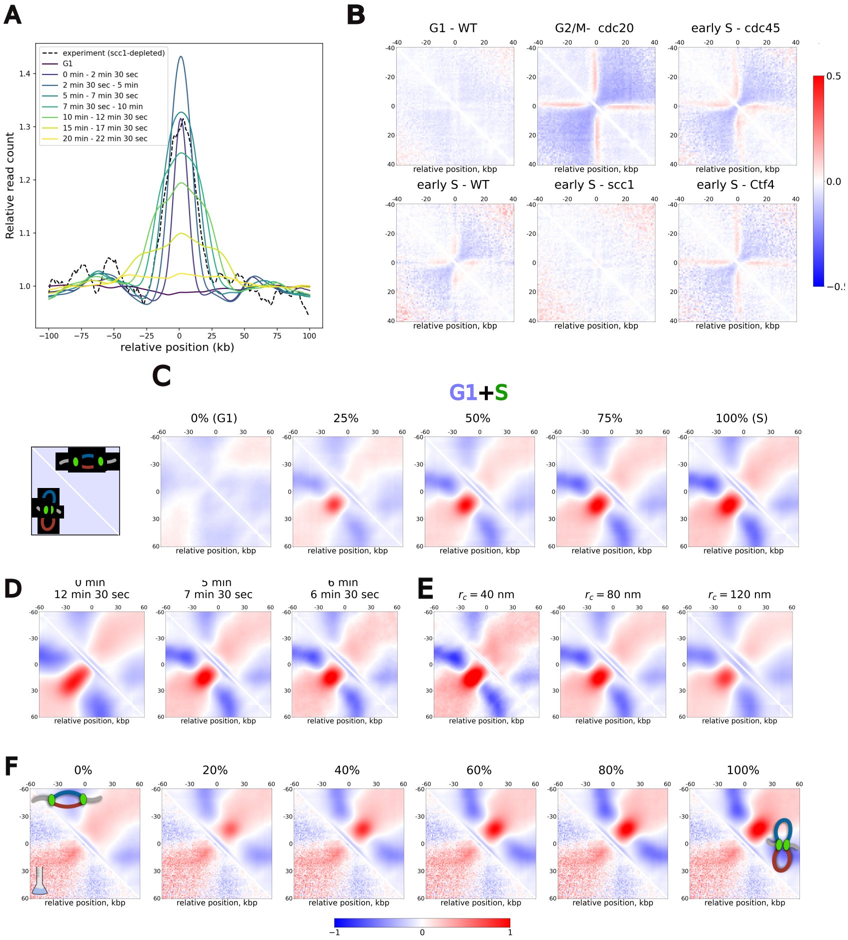

Figure S7: (A) To identify the time in the simulations corresponding to the *in vivo* early-S Hi-C data, we compute the total read count for each genomic position in a 200 kb region around early-replicating origins. Practically, this consists in summing all the contacts of the corresponding rows of the unbalanced contact matrix. To compare experiment and simulations, we divide the signal by the average read count of regions at a distance between 50 to 100 kb from the origin (regions more likely to not being replicated). Analysing the width of the peak (correlated with the typical replicon sizes) we find that times between $t = 5$ and $t = 7$ min and 30 sec give the best match with the experimental signal. (B) Average normalized ($\log_2$ Observed over Expected) contact maps around CARs (see Materials and Methods). The cross-like pattern, signature of CAR-CAR interactions typical of mitotic chromosomes (see cdc20-arrested) starts to be established early-S for all the strains containing cohesin. (*Continue in the next page*)

Figure S7: *(Continued)* This is observed regardless the presence (WT and Ctf4) or absence (Cdc45) of ongoing replication. Despite the only partial depletion of Scc1 (see Materials and Methods), the Scc1-depleted strain does not show significant enrichment of CAR-CAR interactions similar to G1-arrested cells. (C,D,E,F) Average normalized ($\log_2$ Observed over Expected) contact maps around early replicating origins to test possible confounding factors in the enrichment intensity. Unless stated otherwise, to test different conditions, we used as replicating map the one between 5 min and 7 min and 30 sec. Colorbar at the bottom of the image. (C) Dilution of the signal due to Asynchronization. Each map was computed by mixing G1-like, unreplicated trajectories with replicating ones at different percentages and for non-interacting (upper triangle) and interacting (lower triangle) sister-forks. Even with a strong dilution, the replication dependent signals, and their qualitative differences, are still visible. (D) Effect of time interval used to produce HiC maps during replication in the non-interacting (upper triangle) and interacting (lower triangle) sister-fork scenarios. A wider interval, due to the increased heterogeneity in replicon sizes, results into more extended fountains in the presence of sister-forks interactions and to loss of enrichment on the diagonal in their absence. (E) Effects of $r_c$ used to compute contact maps. When $r_c$ increases, we observe a strong loss in the intensity and shape of the enrichment.(F) Mixed configurations of interacting and non-interacting sister-forks. Trajectories from interacting sister-forks simulations are mixed with non-interacting ones with different percentages (upper triangles). The resulting maps are compared with the experimental aggregates for Scc1-depleted cells. Note that to generate the mixed configurations (either between replicating/G1 or between interacting/non-interacting), we always aggregate the raw contact maps first and then re-balance the final matrix.

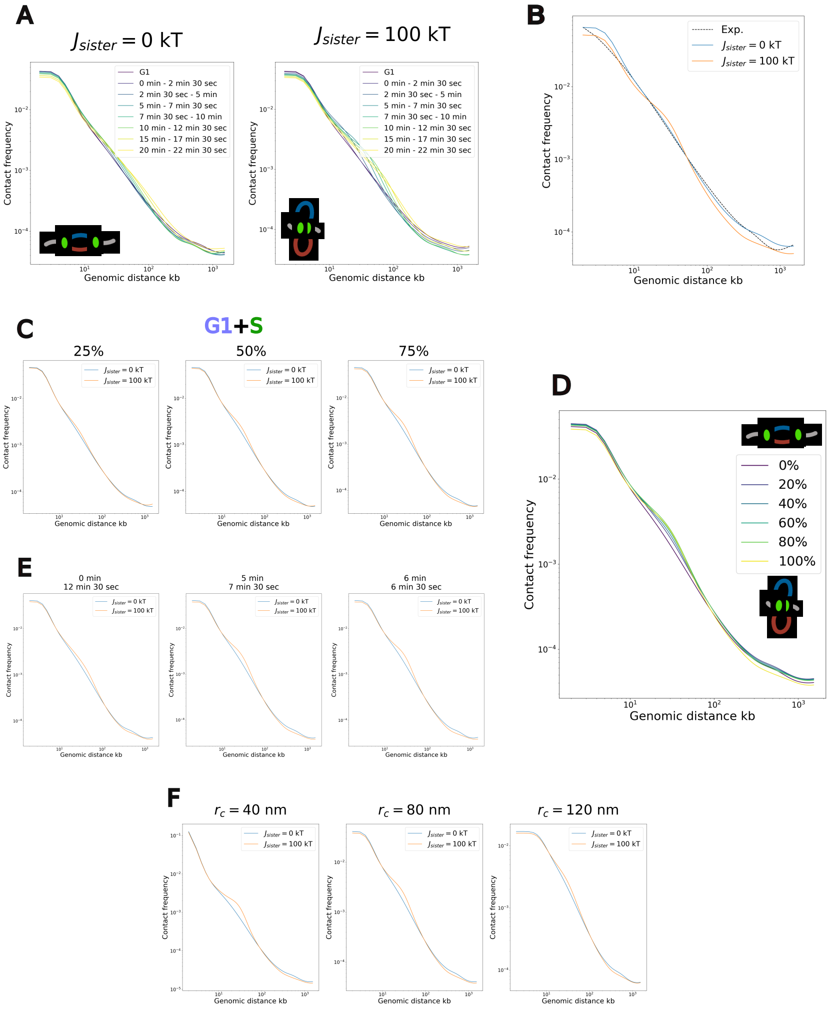


Figure S8: ***P(s)* curves in S-phase.** (A) Time-evolution of *in silico* $P(s)$ in the two scenarios. Maps for interacting sister-forks (Rights) are characterized by a “shoulder”-like feature which evolves according to the replicon size as expected for an extrusion process. (B) Comparison between *in vivo* $P(s)$ (Scc1-depleted cells) and simulations between 5 min and 7 min and 30 sec for non-interacting (blue) and interacting (orange) sister-forks. Simulated curves are rescaled by multiplying by a constant factor $\alpha$ computed as described in the Materialsand Methods ($s_{min} = 10$ kb and $s_{max} = 1$ Mb). Notably, the “shoulder” feature is not clearly observed *in vivo*. *(Continue in the next page)*

Figure S8: *(Continued)* (C,D,E,F) $P(s)$ curves in presence of possible confounding factors as done in Fig. S7 for aggregates around early origins. Unless stated otherwise, to test different conditions, we used as replicating map the one between 5 min and 7 min and 30 sec. (C) Dilution of the signal due to Asynchronization. Each map was computed mixing G1-like, unreplicated trajectories with replicating ones at different percentages and for non-interacting (blue) and interacting (orange) sister-forks. (D) Mixed configurations of interacting and non-interacting sister-forks. Trajectories from interacting sister-forks simulations are mixed with non-interacting ones with different percentages (different curves). (E) Effect of time interval used to produce HiC maps during replication for non-interacting (blue) and interacting (orange) sister-forks. (G) Effects of $r_c$ used to compute contact maps. The computations in (C,D,E,F) show how some of these factors could strongly impact $P(s)$ shape in the interacting case, smoothing the curve towards the non-interacting forks signal. Similarly to Fig. S7, these confounding factor could additively contribute in diluting the shoulder in the *in silico* $P(s)$ and obtain a prediction more consistent to the *in vivo* curve.

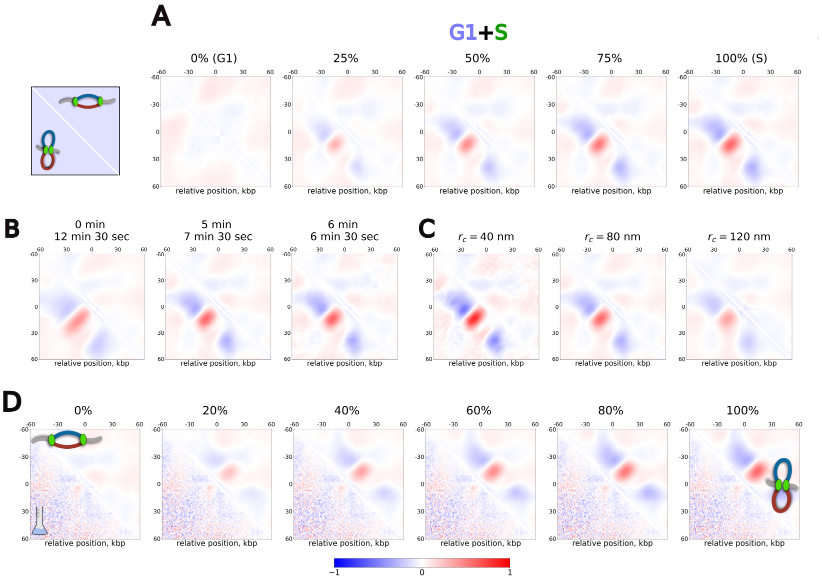


Figure S9: **Aggregate plot around late origins.** (A,B,C,D) Average normalized ($\log_2$ Observed over Expected) contact maps around late replicating origins to test possible confounding factors in the enrichment intensity. Unless stated otherwise, to test different conditions, we used as replicating map the one between 5 min and 7 min and 30 sec. Colorbar at the bottom of the image. All the following results lead to analogous conclusions to the ones described in the context of early replicating origins. (A) Dilution of the signal due to Asynchronization. Each map was computed mixing G1-like, unreplicated trajectories with replicating ones at different percentages and for non-interacting (upper triangle) and interacting (lower triangle) sister-forks. (B) Effect of time interval used to produce HiC maps during replication non-interacting (upper triangle) and interacting (lower triangle) sister-forks. (C) Effects of $r_c$ used to compute contact maps. (D) Mixed configurations of interacting and non-interacting sister-forks. The resulting maps are compared with the experimental aggregates for Scc1-depleted cells.

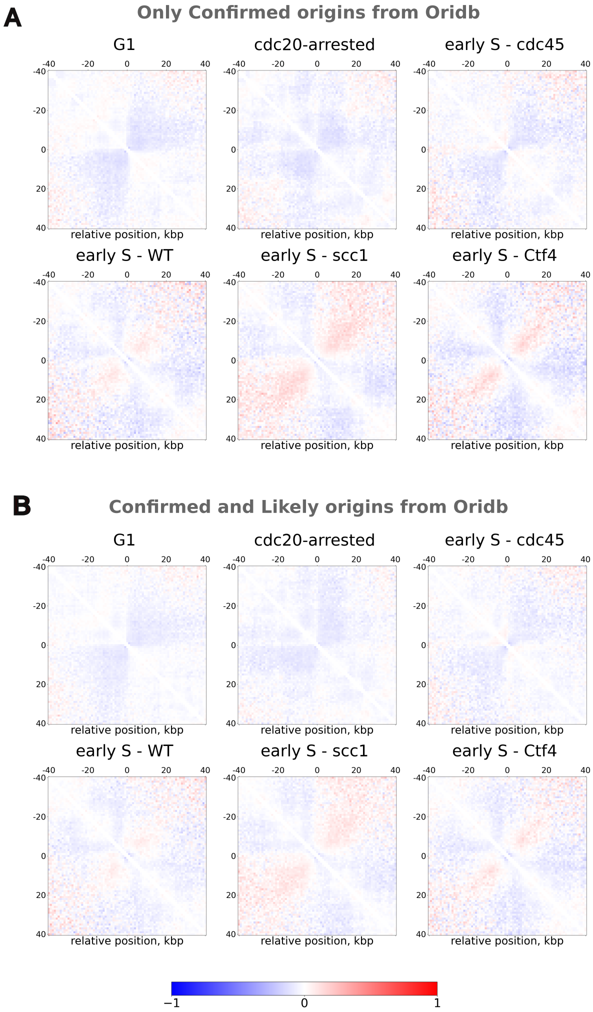


Figure S10: **Aggregate plots around ARS.** Average normalized ($\log_2$ Observed over Expected) *in vivo* contact maps around (A) "Confirmed" (red lines in Fig. S3C) and (B) "Confirmed" and "Likely" ARS (red lines and green in Fig. S3C). The computation was performed for all the mutant strains discussed in the main text with the same conclusion (See Fig. 3A for more information). Consistent with the high number of ARS in the Oridb database which do not correspond to inferred early origins (see Fig. S6C), the signal appears in both cases much lower than Fig. 3A in the main text where, thanks to the predictive power of the $IPLS$, we could restrict the average to origins more likely to be fired in early S-phase.

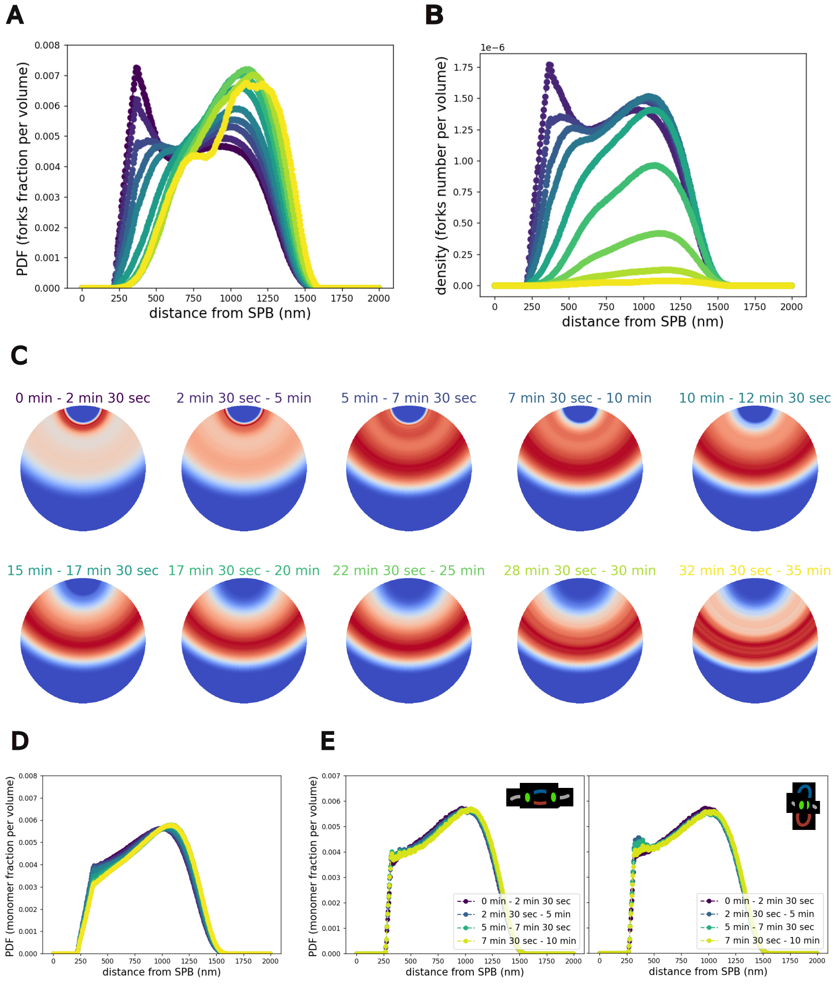


Figure S11: **Spatial distribution of forks in longer simulations**. (A,B) Forks' Probability Distribution Function as a function of the distance $r$ from the SPB (see Materials and Methods) at different time intervals which cover the full S-phase (label of each of the colors can be found in (C). Here, results for the non-interacting sister-forks is presented ($J_{sister} = 0$ kT). On (A), each distribution is normalized to 1 (as in the main text in Fig.4B). On (B) we plot the density of forks without such normalization to highlight the decrease in the forks number at larger $t$. (C) 2D graphical representation of the time evolution of normalized forks densities plotted in (A). This longer set of simulations shows how the remaining forks in late S-phase are enriched on the equatorial plane while the peak in density around the SPB, which is typical of all the DNA in the system (Fig. 4A, Top panels, blue curves) is gradually lost. *(Continue in the next page)*

Figure S11: *(Continued)* (D) Probability Distribution Function to find a monomer at a distance $r$ from the SPB at different time intervals which cover the full S-phase (label of each of the colors can be found in (B). We speculate that due to the density increase, minor changes in the PDF can be observed such as a slight decrease at 250 nm from the SPB (the approximate location of centromere attachment). Note that this effect is mild when compared to the forks' redistribution shown in (A). (E) Monomers' Probability Distribution Function as a function of the distance $r$ from the SPB in presence and absence of sister-forks interactions. We can observe how replication by extrusion leads to a mild enrichment in early S-phase at 250 nm from the SPB relatively to the non-interacting scenario.

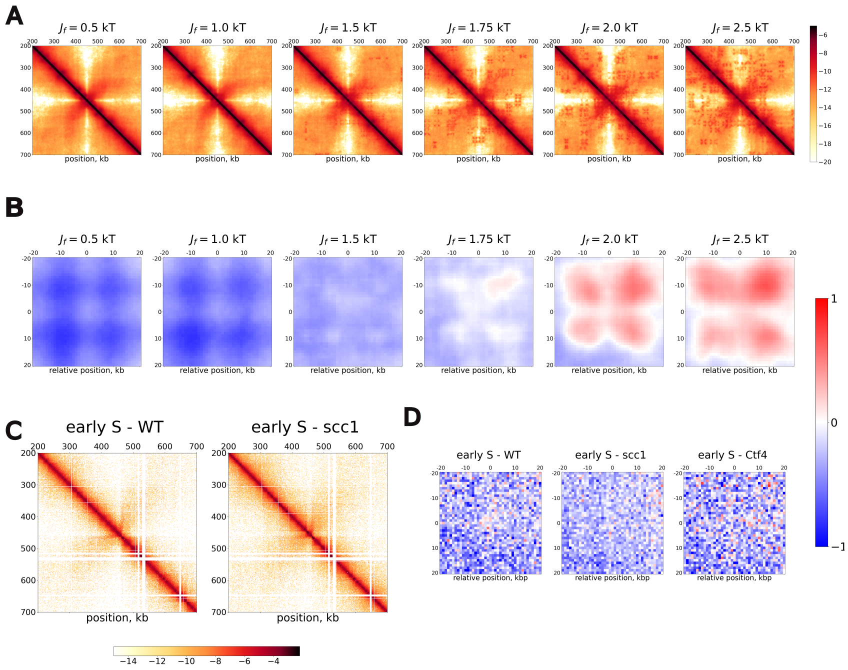


Figure S12: **Model with non-specific aggregative interactions between forks.** (A) Example of normalized HiC maps (500 kb region of chromosome 4) for increasing values of $J_f$ in the interacting sister-fork case ($J_{sister} = 100$ kT). Interestingly, we can clearly observe a transition at $J_f > 1.5$ kT where different forks engage in long range interactions. (B) Such a transition can also be observed in average ($\log_2$ (Observed over Expected)) off-diagonal plots between early origins. For higher $J_f$, we observe a distinct pattern, which reflects the interactions between the 4 forks (4 loops around the inter-origin 0 position). As $J_f$ increases and larger aggregates are formed with higher probability, the intensity increases. *(Continue in the next page)*

Figure S12: *(Continued)* (C) Experimental normalized maps in the same genomic region plotted in (A) of replicating cells in early S-phase (WT and Scc1-depleted). No distinct inter-origin enrichment is detected *in vivo*. (D) Same as (B) for experimental data. On average off-diagonal plots, we do not detect significant inter-origin enrichment in WT or Scc1-depleted cells when compared to the control without replication (early S-phase, Cdc45-depleted).

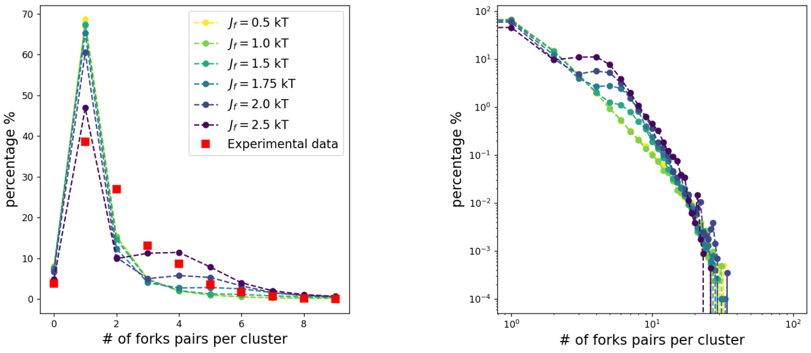


Figure S13: **Detection of RFi for increasing strength of non-specific interaction $J_f$.** Left: percentage of clusters containing a given number of fork pairs as a function of non-specific fork interaction in the interacting sister-fork case ($J_{sister}$ = 100 kT). Red squares indicate the experimental estimate by Saner *et al.*. Simulated data in early-S (between $t = 2$ min and 30 sec and $t = 3$ min and 45 sec) was analyzed to strictly obtain an even number of forks per cluster (see Materials and Methods). Right: Same distributions but in logarithmic scale and including the probability for higher number of forks per cluster. No $J_f$ value seems to drastically improve our comparison. The main effect when increasing energy is a higher probability of stabilizing large clusters.

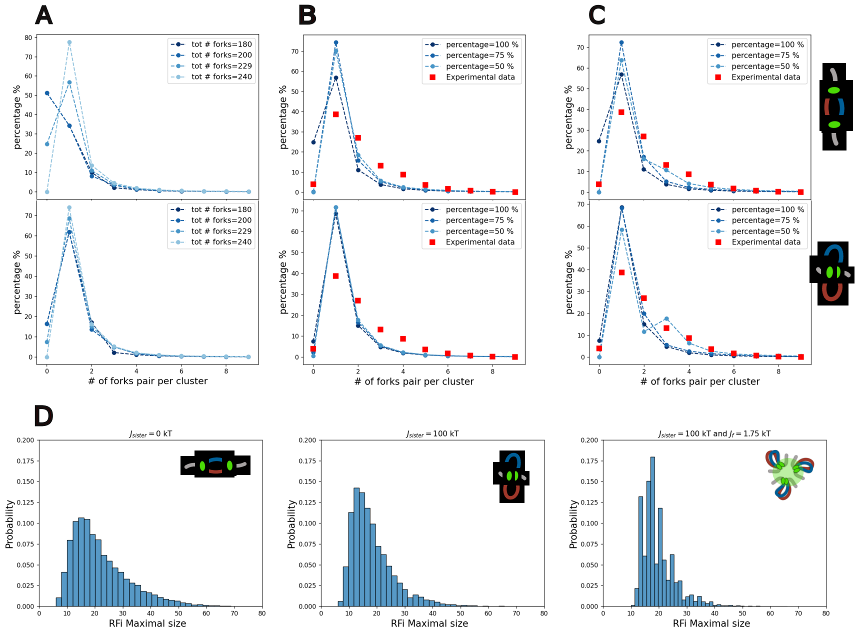


Figure S14: **Possible confounding factors in RFi detection**. As in Fig. 5D in the main text, distributions are computed with a resolution $\theta = 125$ nm. (A,B,C) Top and bottom panels correspond to simulations without and with sister-forks interactions. (A) Effect of the total number of forks used to infer RFi sizes. When compared to the distribution computed with the exact average number of forks (299), we demonstrate how in both scenarios small underestimation or overestimation can quantitatively change the distributions. For example, only 20 more forks are sufficient to eliminate the detection of 0-sized clusters (usually coming from isolated single-forks). (B,C) Effects on the distributions of potentially undetected small clusters. Isolated clusters of size 1(B) or 1 and 2(C) are included in the $A_{i,j}$ association matrix at a given percentage. Again, quantitative differences and increased detection of larger clusters are observed. However, none of these changes can fully account for the inconsistency with the experimental data. (D) Distributions of the largest components (RFi with most forks) of each simulation in the 3 scenarios discussed in the main text (Fig. 5D). Very large clusters containing $\sim 10$ forks are consistently detected. This observation is consistent with the fact that $A_{i,j}$ is commonly characterized by one larger well-connected node and multiple isolated single or sister-forks. For this reason, the probability of having smaller cluster is much higher than intermediate ones (2,3,4 forks pairs), leading to the inconsistency with the experimental estimate.

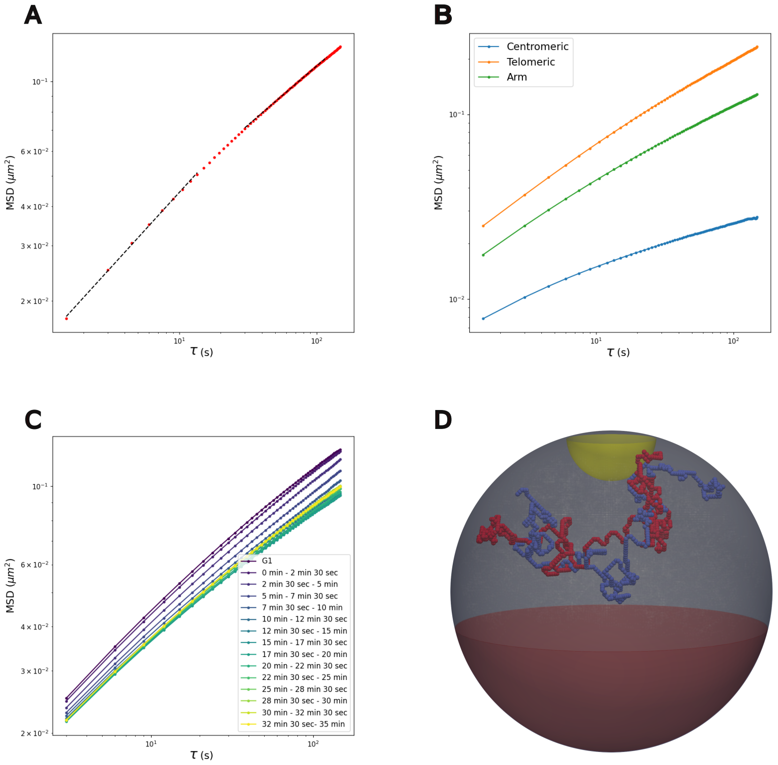


Figure S15: ***MSD* of chromosomes undergoing replication.** (A) Mean Squared Displacement ($MSD$) in G1 (red dots), averaged across the full genome and all the simulated trajectories. The two dashed lines at shorter ($<$ 20 sec) and longer $\tau$ ($>$ 30 sec) indicates the regime where the $MSD$ scales as $\sim$ 0.5 and $\sim$ 0.3 respectively. (B) $MSD$ in G1 for distinct genomic regions: centromeric monomers (distance $<$ 20 kb from the centromere) (blue), telomeric monomers (distance $<$ 20 kb from chains' end) (orange), remaining monomers (green). As expected, centromeres diffuse more slowly due to the strong confinement around SPB. Conversely, monomers in proximity of chain ends (telomeres) diffuse faster. In fact, despite the attachment to the NE, the ends can fluctuate within a distance of 50 nm, likely minimizing its impact on dynamics. *(Continue in next page)*

Figure S15: *(Continued)* (C) $MSD$ for various time windows in S-phase and in the case of non-interacting sister-forks. The curves correspond to the average across the full genome and among all the simulated trajectories. The data correspond to those discussed in Fig. 6A in the main text while including more time-points. (D) Example of Chromosome 8 following replication. The two SCs appear catenated/intertwined as described in our previous study [5].

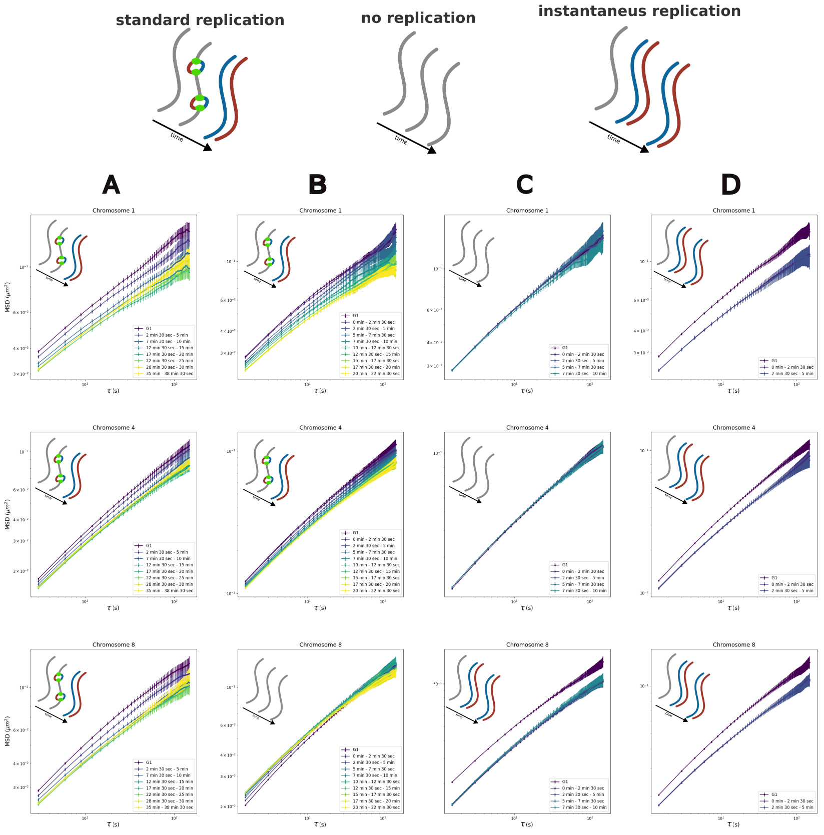


Figure S16: **Effect of replication-driven catenation on** $MSD$**.** Mean Squared Displacement ($MSD$) for different models where normal replication is perturbed measured for individual chromosomes. A short (Chromosomes 1), long (Chromosome 4) and middle (Chromosome 8) chains are selected as examples. (Top) Schemes of the replication status of the chromosomes during the simulation. Standard replication: chromosomes are duplicated according to the regular 1D dynamics described in the main text. No replication: origins assigned to the chromosome cannot be fired and SCs are not synthesized. Instantaneous replication: the two SCs are duplicated very fast (less than one simulation frame) one on top of each other resulting in intertwined structures [5]. *(Continue in the next page)*

Figure S16: (*Continued)* (A) WT-like simulations (Same data used for Fig. S15C and 6C) with standard replication. (B) Model where Chromosome 8 is not replicated (see Materials and Methods). In this scenario the time evolution of density is comparable to the WT case. At large times, only the dynamics of Chromosome 8 (without catenations) is comparable to its G1 counterpart. (C) Model where only Chromosome 8 is instantaneously replicated (see Materials and Methods). In this scenario the overall volumic density remains comparable to the one in G1, while, due to replication-driven catenation, chrosome 8 $MSD$ is reduced. (D) Model where all the genome is instantaneously replicated (see Materials and Methods). For all three chromosomes, we observe a decrease in dynamics which is therefore not dependent on presence the forks in the system.

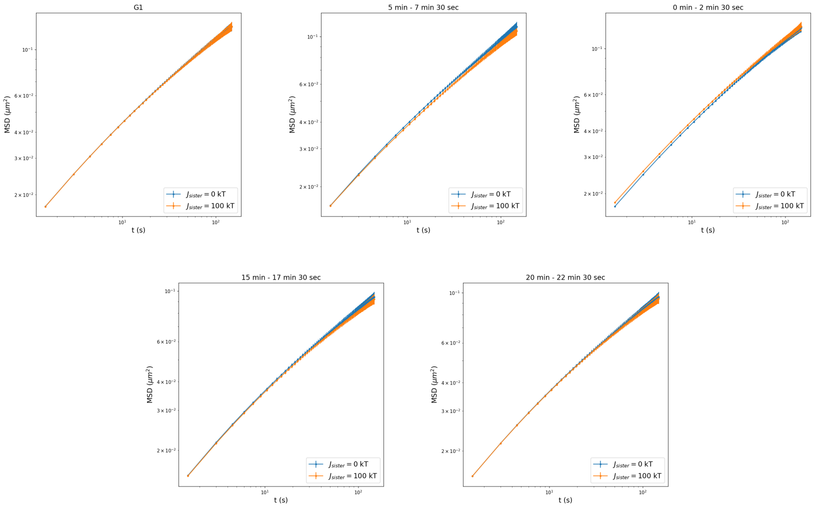


Figure S17: **Comparison of the overall *MSD* in the two models of sister-forks association**. Each panel corresponds to a different time windows used for the computation. The $MSD$ in time does not vary significantly due to the absence (Blue curve) or presence (Orange curve) of sister-forks interactions.

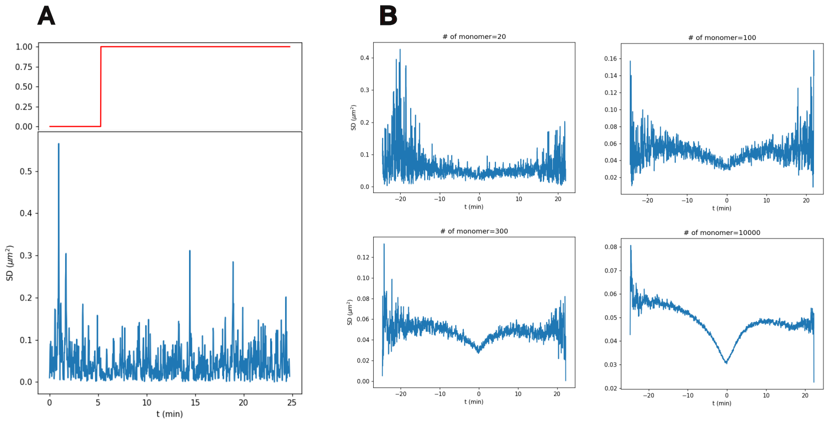


Figure S18: **Squared displacement *SD* around the replication time.** (A) $SD(t, \tau = 15\ sec)$ of a single simulation for an individual monomer. At the top the time evolution of replication status which, around $t = 5$ min goes to 1 (monomer has been duplicated). Fluctuations in $SD$ are too big to observe any effect of replication from a single trajectory. (B) $SD(t, \tau = 15\ sec)$ of a single simulation when $SD$ is averaged among an increasing number of monomers. In particular, a random set of genomic positions were sampled to compute the average. Our analysis suggests that the required number of labeled loci for an individual trajectory (or cells in an experiment) is on the order of 100s.

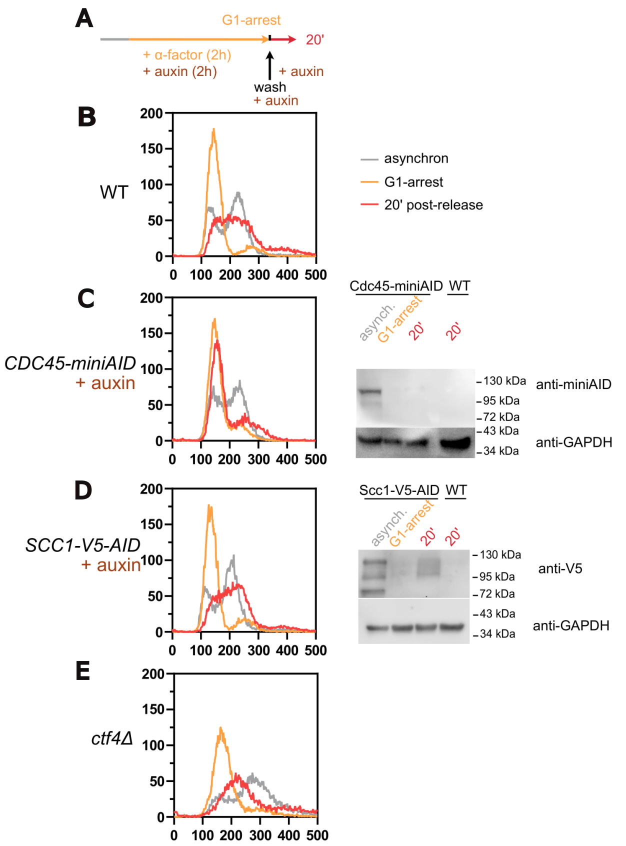


Figure S19: **Cell-cycle progression and protein depletion of experimental Hi-C data.** A) Experimental set-up for G1-synchronisation and release, and for protein depletion. (B,C,D,E) Cell-cycle progression in samples WT (APY607) (B), Cdc45-miniAID (APY539) +auxin (C), Scc1-V5-AID (APY1696) +auxin (D) and Ctf4Δ (APY1814)(E). Western blot are shown in cases of protein depletion (C and D, right). Western blot of Cdc45-miniAID probed with an anti-miniAID antibody and the loading control GAPDH (C, right) and of Scc1-V5-AID probed with an anti-miniAID antibody and the loading control GAPDH (D, right). Expected size of Cdc45-miniAID is 82kDa and expected size of Scc1-V5-AID is 70kDa.

### Supplementary Tables

| Name | # simulations | # processed frames | $t$ between frames |
|---|---|---|---|
| Fig. 1, S1D | 1000 | 400 | $2 \cdot 10^4$ MCS |
| Fig. S2A | 300 | 100 | $2 \cdot 10^4$ MCS |
| Fig. 2B, S2B, S2C, S4, S5 | 500 | 1000 | $4 \cdot 10^4$ MCS |
| Fig. 2E, 3B, 4, 5 | 300 | 100 | $2 \cdot 10^4$ MCS |
| Fig. S7A,C,E,F | 300 | 100 | $2 \cdot 10^4$ MCS |
| Fig. S8A,B,C,D,F | 300 | 100 | $2 \cdot 10^4$ MCS |
| Fig. S9A,C,D, S11E | 300 | 100 | $2 \cdot 10^4$ MCS |
| Fig. S7D, S8E, S9B | 300 | $500, 100, 20$ | $2 \cdot 10^4$ MCS |
| Fig. 5B, S13, S14,S15 A,B | 300 | 100 | $2 \cdot 10^4$ MCS |
| Fig. 6A, S11 A,B,C,D | 100 | 50 | $4 \cdot 10^4$ MCS |
| Fig. S15C | 100 | 50 | $4 \cdot 10^4$ MCS |
| Fig. S12A,B,D, S16 B,C,D | 100 | 100 | $2 \cdot 10^4$ MCS |
| Fig. S12($Jf = 1.75$) | 300 | 100 | $2 \cdot 10^4$ MCS |
| Fig. S16A, S17, S18 | 300 | 100 | $2 \cdot 10^4$ MCS |

Table S1: Statistics of simulated data

**Table S2: Genotypes of *S. cerevisiae* strains used in this study**

| Strains | Genotype | Source |
|---|---|---|
| APY266 | *MATa-inc, ura3::LY-Hocs, lys2::LY, trp1::GAL-HO-hphMX, his3-11,15, RAD5, ade2-1, leu2-3,112, can1-100* | Piazza et al. 2021 |
| RSGY899 | *MATa-inc, ura3::LY-Hocs, lys2::URA3, trp1::GAL-HO-hphMX, his3-11,15, RAD5, ade2-1, leu2-3,112, can1-100, bar1::KAN* | Piazza et al. 2021 |
| APY607 | *MATa-inc, ura3::LY-HOcs, lys2::LY, trp1::GAL-HO-hphMX,* ***his3::OsTIR1-9Myc::HIS3****, RAD5, ade2-1, leu2-3,112, can1::HIS3-LY-Pad* | Piveteau et al. 2025 |
| APY537 | *MATa-inc, ura3::loxP, lys2::URA3, trp1::GAL-HO-hphMX,* ***pMET-CDC20-TRP****, his3-11,15, RAD5, leu2-3,112, ade2-1, can1-100* | Piazza et al. 2021 |
| APY539 | *MATa-inc, ura3::LY-Hocs, lys2::LY, trp1::GAL-HO-hphMX,* ***his3::pADH1-OsTIR1-9Myc::HIS3****, RAD5, ade2-1, leu2-3,112, can1::HIS3-L0.6kb,* ***CDC45-FlagX5-mini-AID::KanMx4*** | Piveteau et al. 2025 |
| APY1696 | *MATa-inc, ura3::LY-HOcs, lys2::LY, trp1::GAL-HO-hphMX,* ***his3::OsTIR1-9Myc::HIS3****, RAD5, ade2-1, leu2-3,112, can1-100,* ***SCC1-Pk3-AID::KMX*** | Piveteau et al. 2025 |
| APY1814 | *MATa-inc, ura3::LY-HOcs, lys2::LY, trp1::GAL-HO-hphMX,* ***his3::OsTIR1-9Myc::HIS3****, RAD5, ade2-1, leu2-3,112, can1::HIS3-LY-Pad,* ***ctf4::KanMX*** | This study |

**Table S3: Summary of Hi-C used in this study**

| Figures | Condition | Strain | Library name(s) | Reference genome | Genome-wide contacts | Source | GEO repository | SRA bioproject | SRA sample |
|---|---|---|---|---|---|---|---|---|---|
| **1, 3A, S1D, S2, S7B, S10** | G1-arrested | APY266, RSGY899 | HB12, HB89 | S288c_R64-2-1 | (50200452 merged), 37291398 | Piazza et al. 2021 | | PRJNA647790 | SRS7056605, SRS9399032 |
| **3A, S7B, S10, S12D** | S-phase | APY607 | AD560 | S288c_R64-2-1 | 18451431 | Piveteau et al. 2025 | | | |
| **3A, S7B, S10** | metaphase-arrest (*cdc20*) | APY537 | AD266 | S288c_R64-2-1 | 28811124 | Dumont et al. 2024 | | PRJNA1139728 | SRX25442218 |
| **3A, S7B, S10, S12D** | S-phase, Cdc45-depleted | APY539 | AD559 | S288c_R64-2-1 | 22451915 | Piveteau et al. 2025 | | | |
| **3A, 3B, S7B, S7F, S8B, S9D, S10, S12D** | S-phase, Scc1-depleted | APY1696 | AD561 | S288c_R64-2-1 | 24834320 | Piveteau et al. 2025 | | | |
| **3A, S7B, S10, S12D** | S-phase, *ctf4Δ* | APY1814 | AD641 | S288c_R64-2-1 | 14210852 | This study | | | |

### Supplementary Videos

- **Video S1**:
  `https://youtu.be/yjj7ZIhEZC0`
  Example of a full genome simulation with chromosomes undergoing replication with sister-forks interactions. Unreplicated DNA is shown with gray beads, while newly synthesized SCs in blue and red. Green beads pinpoint replication forks. The red region within the sphere indicates nucleolus and is inaccessible to other monomers. The yellow shell indicates the surface where centromeres are attached (SPB)

- **Video S2**:
  `https://youtu.be/5VMm6JvkvtY`
  Same as Video S1 but in the case of non-interacting forks and for a longer simulation which cover all S-phase.

- **Video S3**:
  `https://youtu.be/Zggx52Np6OE`
  (Right) Forks in the nucleus in presence sister-fork interactions (same simulation used for Video S1). (Left) Corresponding distribution in function of the distance $r$ from the SPB in time. The distribution is normalized by the corresponding volume of the slice: $V(r+dr) - V(r)$ where $V(r) = (3R - r)r^2\pi/3$. The vertical line corresponds to the position of the yellow spherical shell on the Right.

- **Video S4**:
  `https://youtu.be/ng3g2kbbub8`
  Same as Video S3 but in the case of non-interacting forks and for a longer simulation which cover all S-phase (Video S2).
